# Functional and antigenic constraints on the Nipah virus fusion protein

**DOI:** 10.1101/2025.10.15.682664

**Authors:** Brendan B. Larsen, Sheri Harari, Risako Gen, Cameron Stewart, David Veesler, Jesse D. Bloom

**Affiliations:** Basic Sciences Division and Computational Biology Program, Fred Hutchinson Cancer Center, Seattle, WA 98109, USA; Department of Biochemistry, University of Washington, Seattle, WA 98195, USA; Howard Hughes Medical Institute, Seattle, WA 98195, USA

**Keywords:** Nipah virus, Paramyxovirus, Class I Fusion Protein, Antibody Neutralization, Deep Mutational Scanning, Pseudovirus

## Abstract

Nipah virus is a highly pathogenic virus in the family *Paramyxoviridae* that utilizes two distinct surface glycoproteins to infect cells. The receptor-binding protein (RBP) binds host receptors whereas the fusion protein (F) merges viral and host membranes. Here, we use non-replicative pseudoviruses to safely measure the effects of all F single amino-acid residue mutations on its cell entry function and neutralization by monoclonal antibodies. We compare mutational tolerance in F with previous experimental measurements for RBP and show that F is much more functionally constrained than the RBP. We also identify mutationally intolerant sites on the F trimer surface and core that are critical for proper function, and describe mutations that are candidates for stabilizing F in the prefusion conformation for vaccine design. We quantify how F mutations affect neutralization by six monoclonal antibodies, and show that the magnitude of mutational effects on neutralization varies among antibodies. Our measurements of mutational effects on Nipah virus F predict the ability of the antibodies to neutralize the related Hendra virus. Overall, our work defines the functional and antigenic constraints on the F protein from an important zoonotic virus.

**Importance:** Nipah virus sporadically spills over into humans, where it is often fatal. The Nipah fusion (F) protein is necessary for infection, and is a target for vaccines and antibody therapies. To better understand the constraints on this protein, we experimentally measured how ∼8,500 single amino-acid mutations to F affected its function using pseudoviruses that enable the safe study of protein mutants without the generation of actual replicative virus. We examined the effects of these mutations in the context of structural data and publicly available Nipah virus sequences to characterize the constraints that shape F protein evolution. This work has implications for understanding paramyxovirus fusion proteins, and informs the development of vaccines and monoclonal antibody therapies.

## Introduction

Nipah virus is a bat-borne paramyxovirus that recurrently spills over to humans, causing severe disease with case fatality rates of 40-90% (1–3). Since the first described outbreak in Malaysia in 1998, sporadic spillovers have occurred in India and Bangladesh, with Bangladesh reporting near-annual cases since 2001 (2, 4–6). While these infections typically remain isolated, limited human-to-human transmission has been reported, raising the possibility of wider outbreaks (6, 7).

Nipah virus, like other viruses in the family *Paramyxoviridae*, infects cells through the coordinated action of two different surface glycoproteins: the tetrameric receptor-binding protein (RBP, also known as G) and the trimeric fusion protein (F). F is a class I fusion protein that is cleaved by cathepsins in the endosomal compartment, through a recycling mechanism, to activate it prior to its incorporation into virions (8–11). Once the RBP binds to the host receptors ephrin-B2 or ephrin-B3 (EFNB2/3), it undergoes a conformational change which triggers F to irreversibly transition from its metastable prefusion conformation to an extended postfusion conformation (12–15). During this process, F undergoes large-scale structural rearrangements and inserts an exposed fusion peptide into the host cell membrane, ultimately forming a highly stable six-helix bundle in postfusion F (16–18).

Paramyxovirus RBP tetrameric structures differ substantially, reflecting the wide variety of host receptors used for cell entry (19–23). In contrast, paramyxovirus F proteins show high structural conservation to each other in the pre- and in the postfusion conformations, suggesting shared mechanisms of F function and triggering mechanisms across divergent viruses (17, 24, 25).

Understanding the precise molecular interactions between RBP and F during fusion initiation remains a critical knowledge gap for Nipah virus and other paramyxoviruses. Recently, cryo-EM captured a RBP/F complex from a related paramyxovirus, human parainfluenza virus, which showed one of the RBP heads rests on top of F and likely inhibits the triggering of F until it is released via receptor binding (26). However, there is evidence that the mechanism of F-triggering likely differs among the various paramyxovirus genera (17, 27, 28). For Nipah virus and other members of the genus *Henipavirus*, the precise interaction between RBP and F remains unknown.

Despite the fact that Nipah virus causes severe disease in humans, there are currently no approved vaccines or therapeutics for this virus. Vaccines based on prefusion-stabilized F elicit neutralizing antibodies in mice whereas vaccines based on postfusion F do not (29, 30). Monoclonal antibodies targeting F have shown exceptional promise in preventing disease in animal models, outperforming a best-in-class RBP-directed antibody (31–34). However, evolution can rapidly erode antibody effectiveness for other viruses, sometimes through single mutations which abrogate antibody binding to the viral glycoproteins (35, 36). A better understanding of how mutations affect antibody neutralization can help guide the development of monoclonal antibodies and antibody cocktails with optimized resilience to viral evolution (37).

To provide an in-depth characterization of the function and antigenicity of the Nipah virus F protein, we performed deep mutational scanning on the Nipah F ectodomain using a pseudotyped lentiviral platform that enables us to safely measure the functional and antigenic impact of F mutations without using an authentic Nipah virus isolate. We experimentally measured the effects of all possible single amino-acid residue mutations to Nipah virus F on cell entry and neutralization mediated by six monoclonal antibodies. These data provide detailed information about the functional role of different regions of F, help identify candidate stabilizing mutations for vaccines, and quantify the resilience of different antibodies to possible future viral evolution.

## Results

### Structure and evolution of Nipah virus F

Nipah virus F is a trimeric, type I transmembrane protein made up of three major domains named D1, D2, and D3, which correspond to the lateral, basal, and apical faces, respectively (**Fig. 1A**). In the prefusion structure, heptad repeat A (HRA) forms a series of alpha-helices and beta-sheets which transition into a single long alpha-helix together with the central helix in the postfusion structure (**Fig. 1B**). Heptad repeat B (HRB) forms a three-helix coil leading to the viral membrane in the prefusion F conformation, which refolds to interact with the periphery of the postfusion HRA coiled-coil, leading to the aforementioned six-helix bundle (**Fig. 1B**). An upstream helix is proximal to the central helix, with which it forms a disulfide bond, and that has been shown to have an important role in F triggering (38, 39).

**Figure 1.**
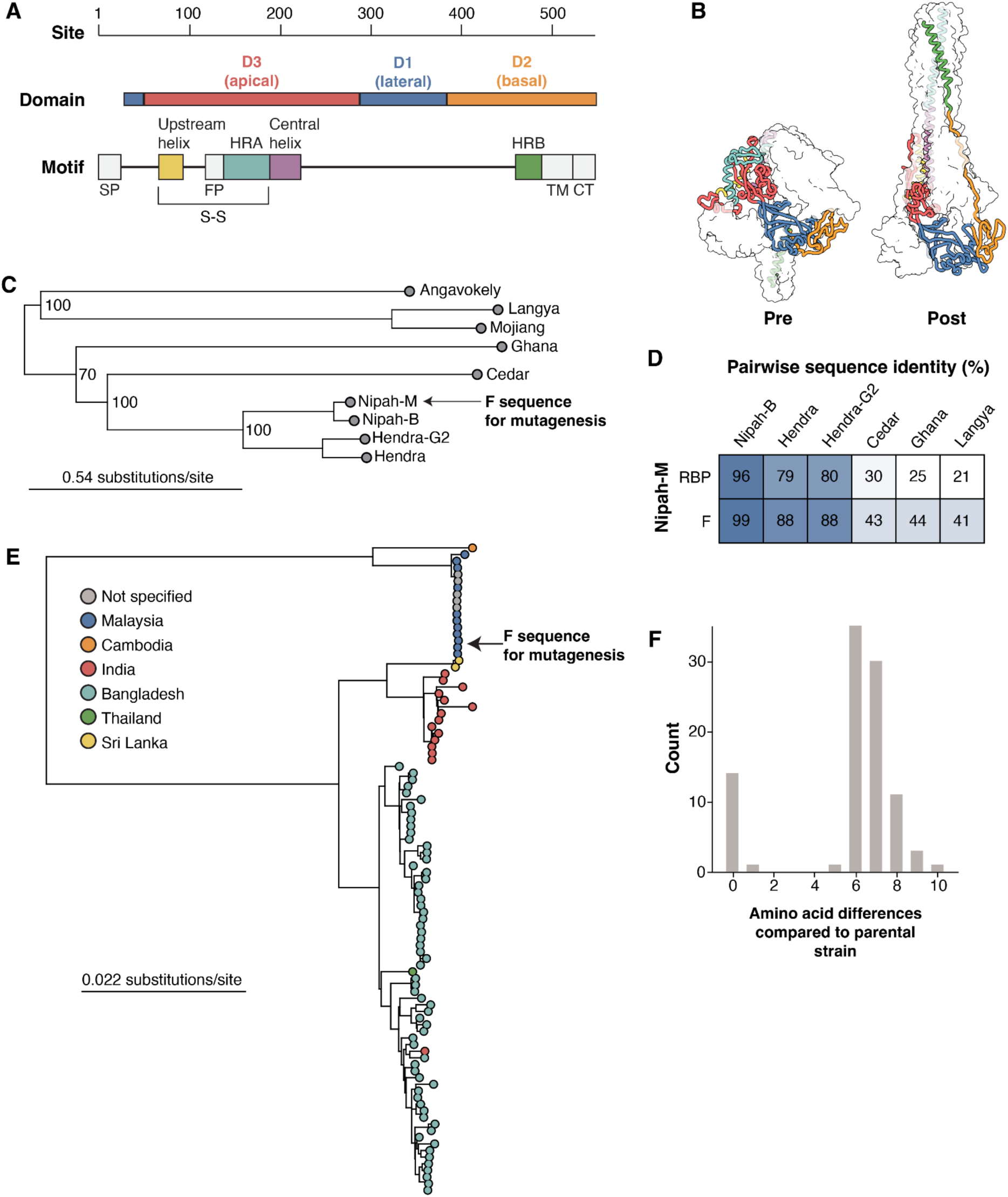
Functional domains and evolution of Nipah F. **A)** Major structural domains and motifs in Nipah F. The motifs are abbreviated as follows. Signal Peptide (SP), Heptad-repeats A and B (HRA,HRB), Fusion peptide (FP), transmembrane region (TM), and cytoplasmic tail (CT). The disulfide bond between F1 and F2 is indicated with a S-S. **B)** Nipah F trimer with one monomer colored by domain or motif as in panel A, and the other two monomers shown as transparent white surfaces. The prefusion structure is based on PDB 5EVM, the postfusion structure is an AlphaFold2-generated structure of postfusion Nipah virus F using the postfusion Langya virus F as template (PDB 8TVE) (21). **C)** Maximum-likelihood phylogenetic tree built from representative publicly available henipavirus whole-genome sequences. Arrow points to Nipah Malaysia, the parental sequence used in the deep mutational scanning experiments. **D)** Pairwise amino-acid sequence identity between Nipah Malaysia RBP and F with other representative henipaviruses. **E)** Maximum-likelihood phylogenetic tree of all Nipah virus whole-genome sequences deposited in GenBank. Circles are colored by the country of origin provided in GenBank records. The arrow points to the Nipah Malaysia strain used as the parent for the deep mutational scanning experiments. **F)** Amino-acid differences in full-length F sequences between the Nipah Malaysia parental strain used in our study and all other Nipah F sequences.

Nipah virus is a member of the *Henipavirus* genus, which includes other viruses that cause disease in humans, such as Hendra virus and the recently described Langya virus (**Fig. 1C**). Across these paramyxoviruses, F is more conserved than RBP at the amino acid level (**Fig. 1D**). For the parental Nipah virus F amino-acid sequence for our deep mutational scan, we used a codon-optimized F gene from the reference strain from GenBank (NC_002728.1), which was sequenced during the first outbreak in 1999 in Malaysia (**Fig. 1E**). Among all publicly available sequences, there is a maximum of ten amino acid differences between the strain used in deep mutational scanning and all other Nipah virus F sequences, despite broad geographical separation and 25 years of evolution (**Fig. 1F**). These data are suggestive of significant functional constraints on Nipah virus F.

### Pseudovirus deep mutational scanning libraries of Nipah virus F

We performed pseudovirus deep mutational scanning using a previously described approach based on pseudotyped lentiviral particles (**Fig. S1**) (40, 41). These pseudoviruses encode no viral proteins other than F and can only undergo a single round of cell entry. They are therefore not fully infectious agents as they do not replicate or cause disease, making them a safe tool to study proteins from deadly pathogens such as Nipah virus. For our experiments, we truncated portions of the cytoplasmic tail of F and RBP, as this increased pseudovirus titers (see previous work (42, 43), Methods, and **Fig. S2**).

To generate the F mutant libraries, we used 300 nucleotide oligo pools (oPools) synthesized by Twist Biosciences that contained all possible amino-acid mutations to the ectodomain of F (site 29 to 481) and assembled them into lentiviral vectors using Golden Gate Assembly (**Fig. S3**), an approach loosely based on previous work (44–46). We added barcodes consisting of 16 random nucleotides downstream from the stop codon of each F variant; these barcodes are subsequently linked to the full F sequence by long-read PacBio sequencing which then facilitates downstream sequencing by enabling each variant to be identified by Illumina sequencing of its barcode (40).

We made two replicate libraries (LibA and LibB), which have unique mutation-barcode linkages, followed by generation of cell-stored lentiviral libraries in 293T-rtTA cells, where each cell contains a single provirus with a unique Nipah F variant (40, 41). There were 56,368 and 62,614 unique barcoded variants for LibA and LibB, respectively. 69% of these variants contained a single amino-acid mutation relative to the parental sequence, with the remainder being unmutated (∼11%) or variants with >1 mutation (∼20%) (**Fig. S4A**). Most mutations relative to the parental sequence matched the exact codon sequence used for the oPool design (**Figs. S4B, S4C**). Most of the variants with >1 mutation likely arose from lentiviral recombination (47, 48), an unavoidable step in generating our pseudovirus libraries (**Fig. S4D**). These results demonstrate low error rates in the oPool synthesis and library assembly.

### Measuring effects of Nipah F mutations on cell entry

To measure the effects of F mutations on cell entry, we used an approach similar to that recently described in our deep mutational scanning of Nipah RBP (41). Specifically, we quantified the ability of pseudoviruses with each F protein mutant to enter a cell line expressing a Nipah virus receptor protein relative to control pseudoviruses that use a different viral entry protein (VSV-G) to enter cells. In these experiments, mutations to F can affect cell entry through several different processes, including by affecting protein expression, folding, stability, and triggering by RBP. Mutation effects are reported as log_2_ transformed values, where −1 represents a two-fold reduction in cell entry relative to the parental unmutated F protein. Since ∼20% of the variants contain multiple mutations (**Fig. S4A**), we fit global epistasis models to estimate the effects of single mutations from the entire library of single and multiple mutant variants (49). The effects of mutations determined from the global epistasis model fitting were highly correlated with the effects of mutations in the single-mutant variants only (see Methods, **Fig. S5**).

Nipah virus can use two different receptors, ephrin-B2 or -B3 (EFNB2/3), to enter cells (15, 50, 51). We previously generated CHO cells that stably express either *Pteropus alecto* bat ephrin-B2 or -B3 (CHO-bEFNB2/3) (41). The use of cells expressing bat rather than human ephrin receptors reduces the possibility that our experiments could uncover potentially hazardous information about F protein mutations that might adapt the virus to better infect human cells (41, 52, 53). In prior work, we found that the effects of mutations on the cell entry function of Nipah RBP differ somewhat between cells expressing bEFNB2 versus bEFNB3 (41). Although F is not directly involved in receptor binding, prior work has suggested that F is triggered more effectively by RBP binding to EFNB2 relative to EFNB3, possibly because RBP binds to EFNB2 more strongly (54). However, we found the average effects of F mutations at each site on cell entry were highly correlated between measurements made on bEFNB2 versus bEFNB3 expressing cells, in sharp contrast to similar measurements for RBP mutations (r = 0.99 for F versus 0.82 for RBP; **Fig. S6**). Therefore, at least in our experimental system, the effects of F mutations on cell entry are largely independent of whether the target cells express bEFNB2 or bEFNB3, a result that makes sense in light of F’s primary function being membrane fusion rather than receptor binding. For all subsequent results in this paper, we report the effects of mutations from experiments performed in CHO-bEFNB3 cells.

In total, we reliably measured the effects of 8,449 F mutations on cell entry, which represents 98.2% of all possible amino-acid mutations. Each mutation had an average of 5.9 and 6.2 associated unique barcodes for LibA and LibB, respectively (**Fig. S7A**). The effects of mutations on entry were highly correlated between our replicate variant libraries (r=0.94; **Fig. S7B**), and throughout the rest of this paper we report the average of the measured effects across libraries.

### Effects of Nipah F mutations on cell entry

Examination of our comprehensive measurements of mutational effects on cell entry shows that the F protein is highly mutationally constrained (**Fig. 2**), which is consistent with the low rate of protein evolution observed in circulating sequences (**Figs. 1D, 1F**). We compared the effects of mutations on entry with our previously generated RBP data (41). In general, F mutations were much more deleterious for cell entry compared to RBP mutations (**Fig. 3A**). This result indicates the lower rate of natural sequence evolution in F relative to RBP is a result of higher functional constraints on the protein.

**Figure 2.**
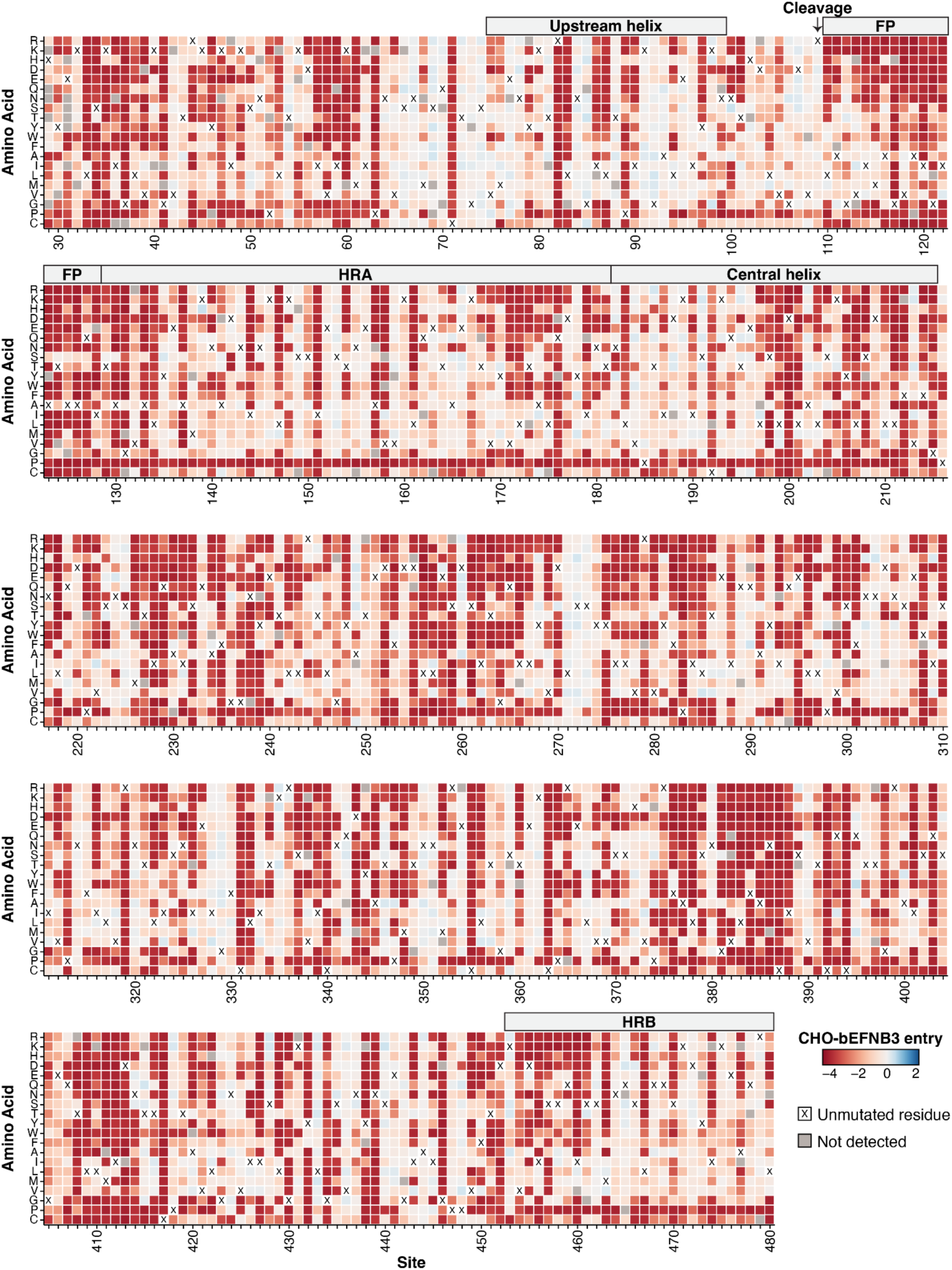
Effects of Nipah F mutations on entry in CHO-bEFNB3 cells. Each mutation is colored by its effect on cell entry. Effects on cell entry range from highly deleterious (red) to neutral (white) to slightly beneficial (blue). The amino acid identity at each site in the parental F from the Malaysia strain is indicated with a black ‘X’. Mutations that were missing in our library and so lack a measurement are colored gray. The experiments were performed in CHO cells that stably express ephrin-B3 from the bat species *Pteropus alecto*. See https://dms-vep.org/Nipah_Malaysia_F_DMS/visualizations/posts/CellEntryHeatmapWrappedZoomable.html for an interactive version of the plot.

**Figure 3.**
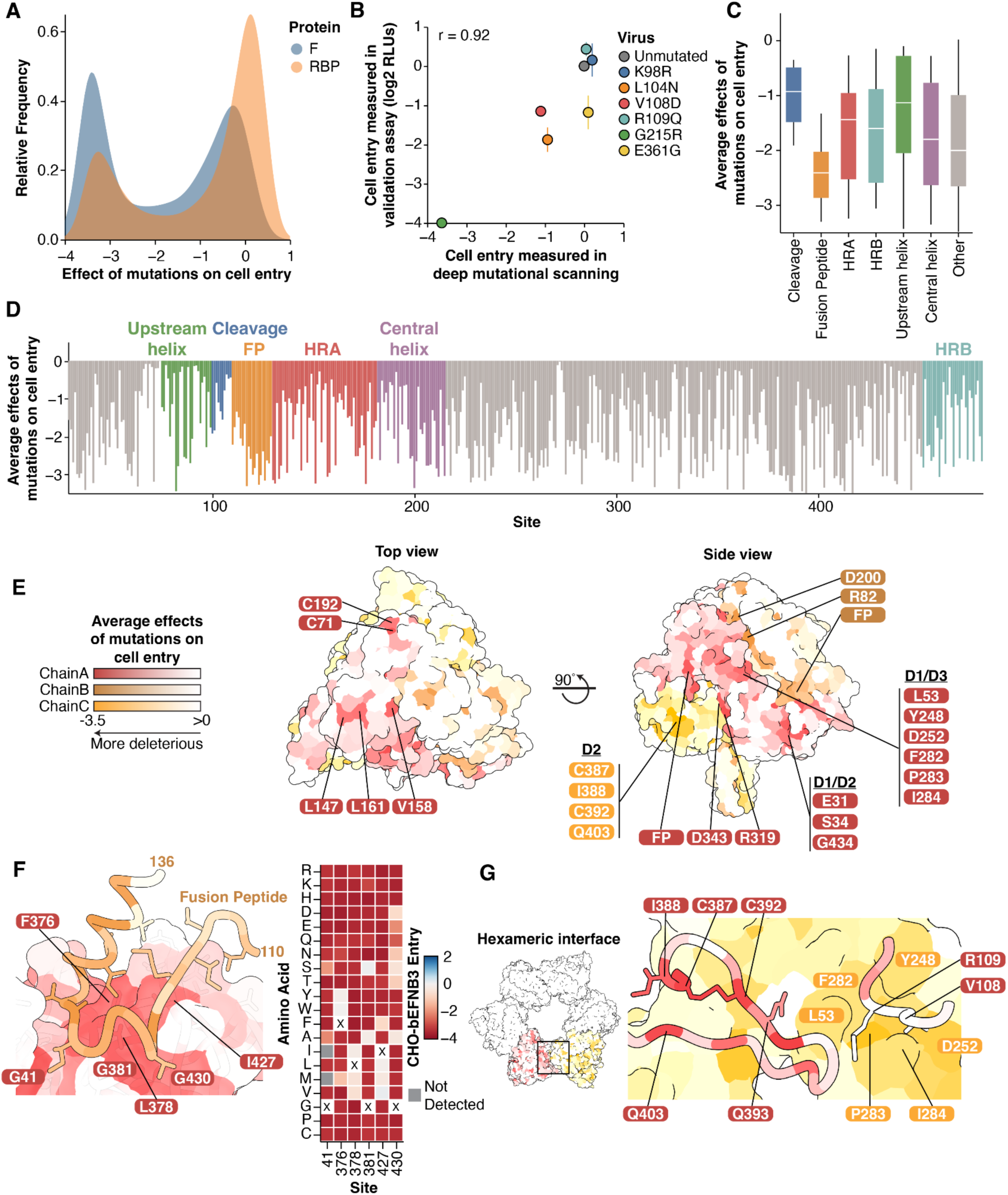
Effects of F mutations on cell entry in context of the protein structure. **A)** Distribution of the effects of all amino-acid mutations to F or RBP on cell entry in CHO-bEFNB3 cells, with the measurements for F from the current study and RBP from Larsen et al. 2024 (41). More negative values indicate mutations more strongly impair cell entry. **B)** Validation of cell entry effects in CHO-bEFNB3 cells. Correlation between deep mutational scanning cell entry measurements and individual pseudoviruses expressing the respective mutation. Individual mutations were chosen to span a range of entry effects. For validation assays, relative light units (RLUs) were measured by a luciferase assay and compared to the unmutated values. Measurements were done in triplicate. **C)** Min-max boxplot showing the average effects of all amino-acid mutations at each site for different regions of F. **D)** Average effect of mutations at each site across the F ectodomain, colored by protein domain or region. **E)** Average effects of mutations at each site mapped onto the prefusion F structure (PDB: 5EVM), viewed from the top and side. Highly constrained, surface-exposed residues are labeled. Each protomer is colored with a separate color. **F)** The fusion peptide rests on a highly constrained surface of the nearby protomer in the prefusion F structure. Sites are colored by the average effects of mutations on cell entry. Key constrained interface residues are labeled. The heatmap (right) shows the effects of all mutations at the sites labeled on chain A on the structure. **G)** Nipah virus F trimers are arranged in a hexameric ring, as observed in the crystal lattice described in PDB: 5EVM. The boxed area at the interface of the trimers is shown in more detail on the right. Trimers are colored separately. Specific residues at the interface are labeled.

To validate our deep mutational scanning measurements of the effects of F mutations on cell entry, we generated pseudoviruses expressing different individual F mutations selected to span a range of effects, and compared their entry relative to the unmutated parental strain. Entry into cells for the individual pseudovirus mutants was highly correlated with the deep mutational scanning measurements of the effects of these mutations (r=0.92; **Fig. 3B**).

Based on the average effects of F mutations at each site on cell entry, the fusion peptide is the most constrained region, while the region upstream of the cleavage site is fairly tolerant to mutations (**Figs. 3C, 3D**). These data are congruent with previous low-throughput mutagenesis experiments on Nipah F within the cleavage and fusion peptide regions (10, 55–57). Of the four helical regions, the upstream helix was the least constrained (**Fig. 3C**).

To understand the structural context of mutational constraint, we mapped the average effects of mutations at each site onto the trimeric prefusion structure (**Fig. 3E**). The surface-exposed apex of the protein is largely tolerant to mutations, with the exception of two cysteine residues which form a disulfide bond between F1 and F2 (C71/C192) and three hydrophobic residues (L147, V158, and L161; **Fig. 3E**).

The surface-exposed lateral face of the trimer is more constrained than the apex. Some of the most highly constrained surface-exposed sites are charged residues, including R82, D200, R319, and D343 (**Fig. 3E**). There are five additional patches of surface-exposed residues on the lateral faces that are highly constrained. These include the fusion peptide interface between the monomers, a region consisting exclusively of residues in D2 (C387, I388, C392, Q403), a region of D1/D2 (E31, S34 G434), and a region in D1/D3 (L53, Y248, D252, F282, P283, I284) (**Fig. 3E**).

The fusion peptide (sites 110-128) is inserted into a groove in the opposing protomer (**Fig. 3F**). Fusion peptide residues directly in this interface were more constrained than those outside of it (**Fig. 3F**). The cavity-forming residues surrounding the fusion peptide are primarily composed of hydrophobic and glycine residues, and the effects of mutations at these sites are defined by specific amino acid biochemical properties. For example, at sites G41 and G381, mutations to amino acids with small side chains (A,S) were less deleterious than other mutations (**Fig. 3F**). These data suggest that efficient sequestration of the fusion peptide is critical for proper F function.

Prior work has suggested that Nipah virus F trimers may assemble into hexameric rings (25, 58). Two of the surface-exposed patches of constrained residues (D2 and D1/D3; **Fig. 3G**) are within the interface between trimers of the hexamer (**Fig. 3G**). In one trimer, the loop containing the cleavage site at R109 contacts the opposing trimer at sites in the highly constrained D1/D3 patch, including L53, P283, and I284. The constrained patch from D2, including C387, I388, C392, and Q403 also contact the opposing trimer (**Fig. 3G**). While the cysteine residues are important for maintaining overall protein structure, these data are consistent with an important functional role of the hexameric interfaces in modulating fusion (58).

### Effects of mutations mapped to the postfusion structure

Since Nipah virus F transitions from a prefusion to a postfusion conformation during viral entry, we next examined the effects of mutations in the postfusion structure. Some highly constrained residues that were buried in the prefusion structure have moved to the surface (**Fig. 4A**), but the average effects of surface exposed residues remains similar to the prefusion structure (**Fig. 4B**).

**Figure 4.**
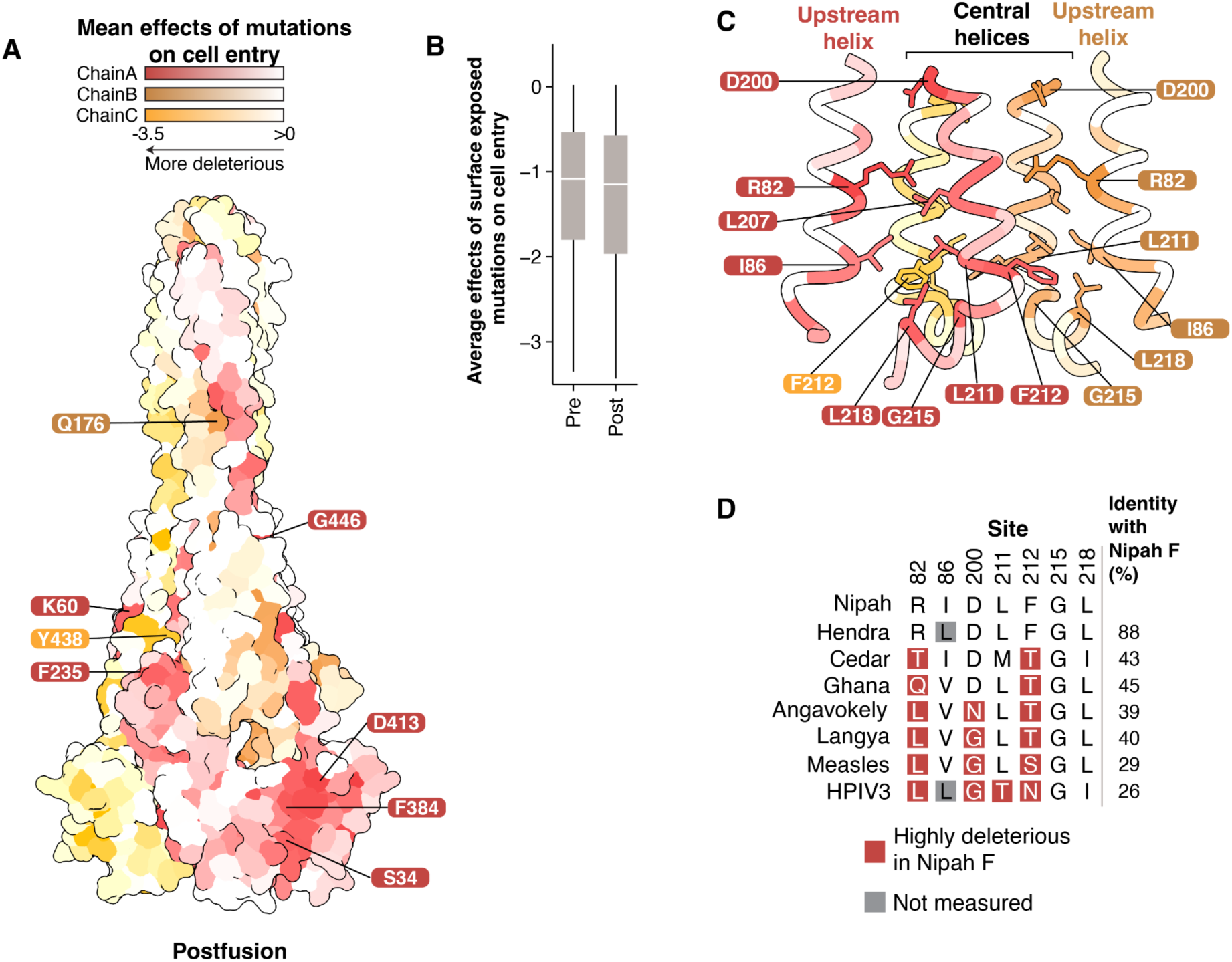
Effects of mutations on cell entry mapped on the postfusion conformation of the F protein. **A)** Average effects of mutations at each site on cell entry mapped onto an AlphaFold2-generated structure of postfusion Nipah virus F using the postfusion Langya virus F as template (PDB 8TVE) (21). Highly constrained surface-exposed sites that were buried in the prefusion structure are labeled. **B)** Effects of mutations at all surface-exposed sites in the pre- and postfusion conformations. Boxed regions show the mean, Q1, and Q3. **C)** Average effects of mutations at each site mapped onto a region near the base of the central helix in the postfusion conformation. R82 in the upstream helix is pointed inward towards the central helices. F212 is buried in a hydrophobic patch formed by I86, L211, G215, and L218. **D)** Amino-acid identity of representative paramyxoviruses at constrained sites labeled in (B). Mutations at sites R82, D200, L211, and F212 that are highly deleterious for cell entry of Nipah F are present as the wildtype amino acid in F proteins of some more divergent paramyxoviruses. Overall pairwise amino-acid identity to Nipah F is shown on the right.

One of the major structural rearrangements is located in D3 and is associated with the extension of HRA into a single helix continuous with the central helix, and the packing of the three HRA helices as a coiled-coil in the center of the trimer (59). To identify key residues involved in this transition, we examined the base of the central helix and the associated upstream helix in the postfusion structure. This region contains many sites that are highly constrained, including R82, D200, F212, and G215 (**Fig. 4C**). R82 and D200 are the most mutationally intolerant amino acids in the upstream and central helices, respectively (**Fig. S8**), and in the prefusion structure they are partially surface-exposed (**Fig. 3E**). However, in the postfusion structure, both R82 and D200 are buried, near the core of the trimer (**Fig. 4C**). F212 is buried in a hydrophobic pocket that includes sites I86, L211, and L218 (**Fig. 4C**). Despite the high constraint observed at these sites for Nipah virus F, the F proteins of other highly divergent paramyxoviruses with <50% amino-acid identity with Nipah F contain amino acids at sites 82, 200, 211, and 212 that are highly deleterious for cell entry in Nipah virus F (**Fig. 4D**). This finding suggests the effects of specific F mutations on cell entry can differ in highly divergent paramyxoviruses, likely due to epistasis with other changes that alter sequence-function constraints.

### Identification of candidate mutations for stabilizing F in the prefusion conformation

For many viral vaccines, adding mutations to the fusion protein that stabilize its prefusion conformation can increase the vaccine-elicited neutralizing antibody titers (60, 61). Several prior studies have described mutations that stabilize Nipah or Hendra F in the prefusion conformation (21, 29, 30, 62), but the identification of additional stabilizing mutations could further improve immunogen design. One previously described Nipah virus F stabilized construct consisted of two cysteine mutations to form an additional disulfide bond between F1 and F2 (L104C/I114C), a space-filling mutation (L172F), and a proline mutation in the central helix (S191P) (29).

The transition of F to the postfusion conformation is necessary for it to mediate cell entry, so we predict prefusion-stabilizing F mutations should be deleterious for cell entry in our deep mutational scanning. As expected, the two previously described mutations that individually stabilize prefusion F (L172F and S191P) were deleterious for cell entry in our measurements (**Fig. 5A**). Two previously described cysteine mutations (L104C and I114C) that stabilize prefusion F by forming a disulfide bond were relatively well tolerated for cell entry in our measurements (**Fig. 5A**), likely because we only measured the impact of single mutations and both cysteine mutations must be made simultaneously to form the stabilizing disulfide bond.

**Figure 5.**
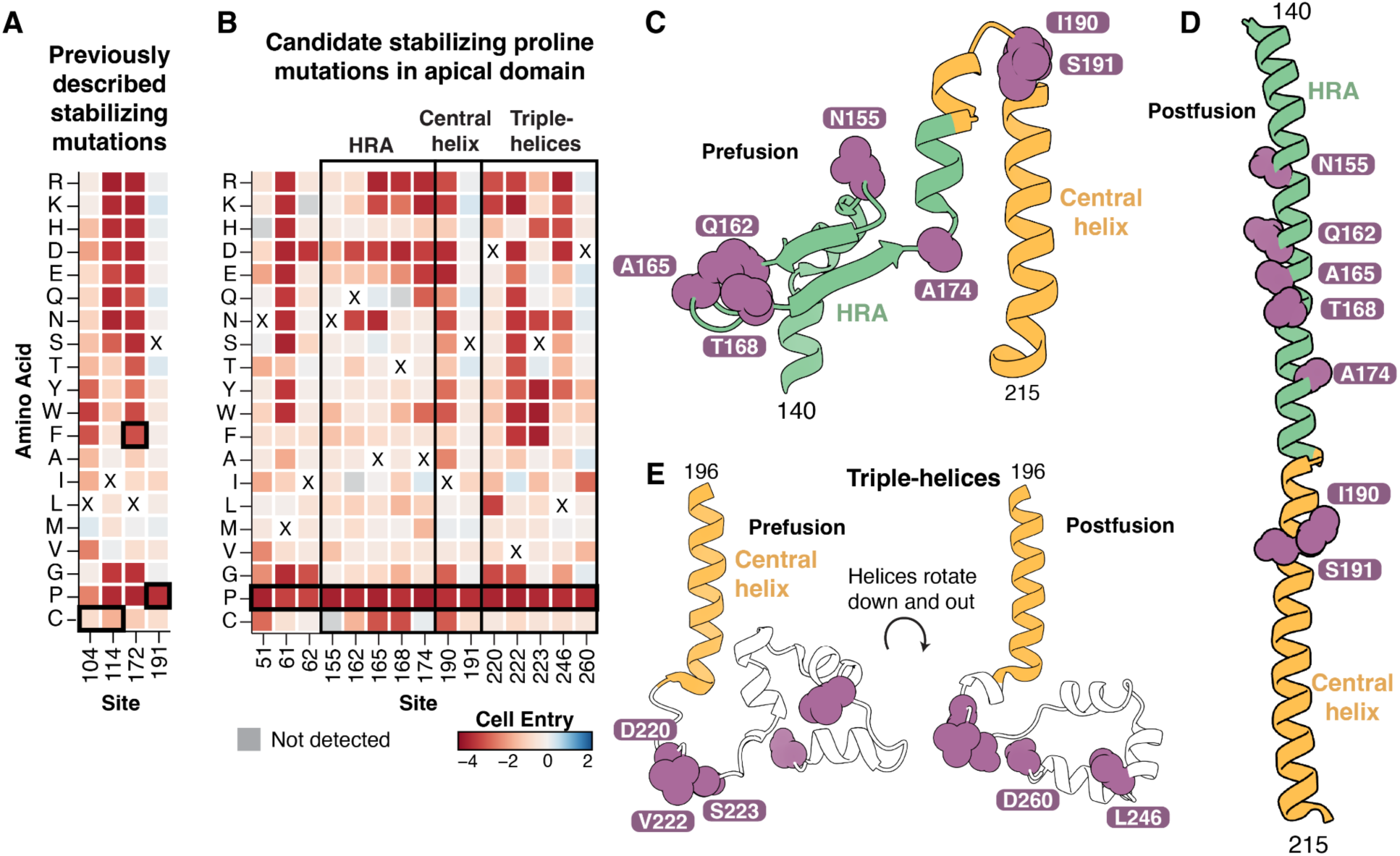
Candidate prefusion stabilizing mutations. **A)** Effects of single mutations on cell entry at sites with previously described (29) prefusion stabilizing mutations (L104C/I114C, L172F, and S191P; each surrounded by a black box). The two cysteine mutations create an additional disulfide bond between F1 and F2, L172F mutation is space-filling, and S191P adds a proline that disrupts helical formation in the postfusion conformation. These mutations are likely deleterious for cell entry even though they increase prefusion F stability because they restrict conformational transitions necessary for F-mediated fusion. **B)** Effects of single mutations on cell entry at sites in the apical domain where we suggest that proline mutations might increase the stability of prefusion F. Sites with highly deleterious proline mutations, and at least some tolerated mutations, likely disrupt helical formation or conformational flexibility, restricting the transition to the postfusion conformation. Heavy black box is around the candidate proline mutations, the light black box surrounds different regions of the apical domain. **C)** Sites with candidate stabilizing mutations (purple) mapped onto HRA (green) and the central helix (yellow) from the prefusion structure. **D)** Sites with candidate stabilizing mutations mapped onto the HRA and central helix from postfusion structure. **E)** Sites with candidate stabilizing mutations in the triple-helix region downstream from the base of the central helix mapped onto the pre- and postfusion structures.

To identify additional candidate prefusion stabilizing mutations to Nipah virus F, we searched for sites where mutations to prolines were highly deleterious, but mutations to most other amino acids were well tolerated, similar to the known stabilizing mutation S191P (**Fig. 5A**). Prolines are uniquely unfavorable in the helical conformations that are adopted in the postfusion conformation, so sites that tolerate mutations to other amino acids but not proline will often be ones where folding of the prefusion conformation is tolerant of mutations but proline specifically impairs cell entry by preventing the transition to the postfusion conformation. We focused on sites in the apical domain since it undergoes a large conformational shift that results in a long alpha-helix in the postfusion conformation (59), and excluded sites that were in helices or strands in the prefusion conformation because proline mutations in these secondary-structure elements are likely to cause misfolding of the prefusion conformation. We identified 15 sites where prolines were highly deleterious but at least four other mutations were tolerated (**Fig. 5B**); five of these sites were in HRA and two were in the central helix (**Figs. 5C,D**), while five were in the triple-helix region just downstream from the central helix base (**Fig. 5E**). Overall, these mutations represent promising candidates for further investigation as prefusion-stabilizing F mutations for vaccine design, although further experimental characterization beyond the scope of the current study would be needed to confirm which of these mutations actually effectively stabilize prefusion F and retain native antigenicity.

### Effects on cell entry of mutations found in circulating Nipah and Hendra viruses

We collated all mutations in the F protein from publicly available Nipah virus sequences and examined their effects on cell entry as measured in our experiments. Nearly all naturally occurring Nipah virus mutations in the regions we mutagenized (the F ectodomain) were neutral (**Fig. S9**). We then took the same approach with available Hendra virus F sequences, which share ∼88% amino-acid identity with Nipah F. If epistasis with other mutations is modifying the effects of mutations in Hendra virus F versus Nipah virus F, we would expect some naturally occurring Hendra virus mutations to be deleterious for cell entry in Nipah virus F. Instead, nearly all mutations in Hendra virus F relative to the Nipah virus F parental strain were also neutral for cell entry (**Fig. S9**).

### Measurement of how all mutations affect neutralization quantifies antibodies resilience to F-protein mutations

We measured how all functionally tolerated mutations to Nipah virus F affect neutralization by six monoclonal antibodies that target the apical (12B2, 2D3, 4H3), lateral (1A9, 1F2), and basal (2B12) faces (33, 63). To measure how F mutations affect antibody neutralization, we performed selections on our replicate pseudovirus libraries with varying concentrations of antibody as previously described (40, 41). Briefly, we quantify infection by pseudoviruses with each F protein mutant relative to a standard that is unaffected by anti-F antibodies in order to measure the impact of each mutation on neutralization.

The mutations that affected neutralization by the antibodies were almost entirely surface exposed and directly mapped to the structurally determined epitopes of each antibody (**Fig. 6A**). To validate the ability of deep mutational scanning to accurately predict neutralization measurements, we generated pseudoviruses expressing different F mutations spanning a range of effects, and measured their neutralization with antibody 4H3. The IC_50_ estimates against the different F mutants as measured in these pseudovirus neutralization assays were highly correlated with the deep mutational scanning data (r=0.96, **Figs. 6B, S11A**).

**Figure 6.**
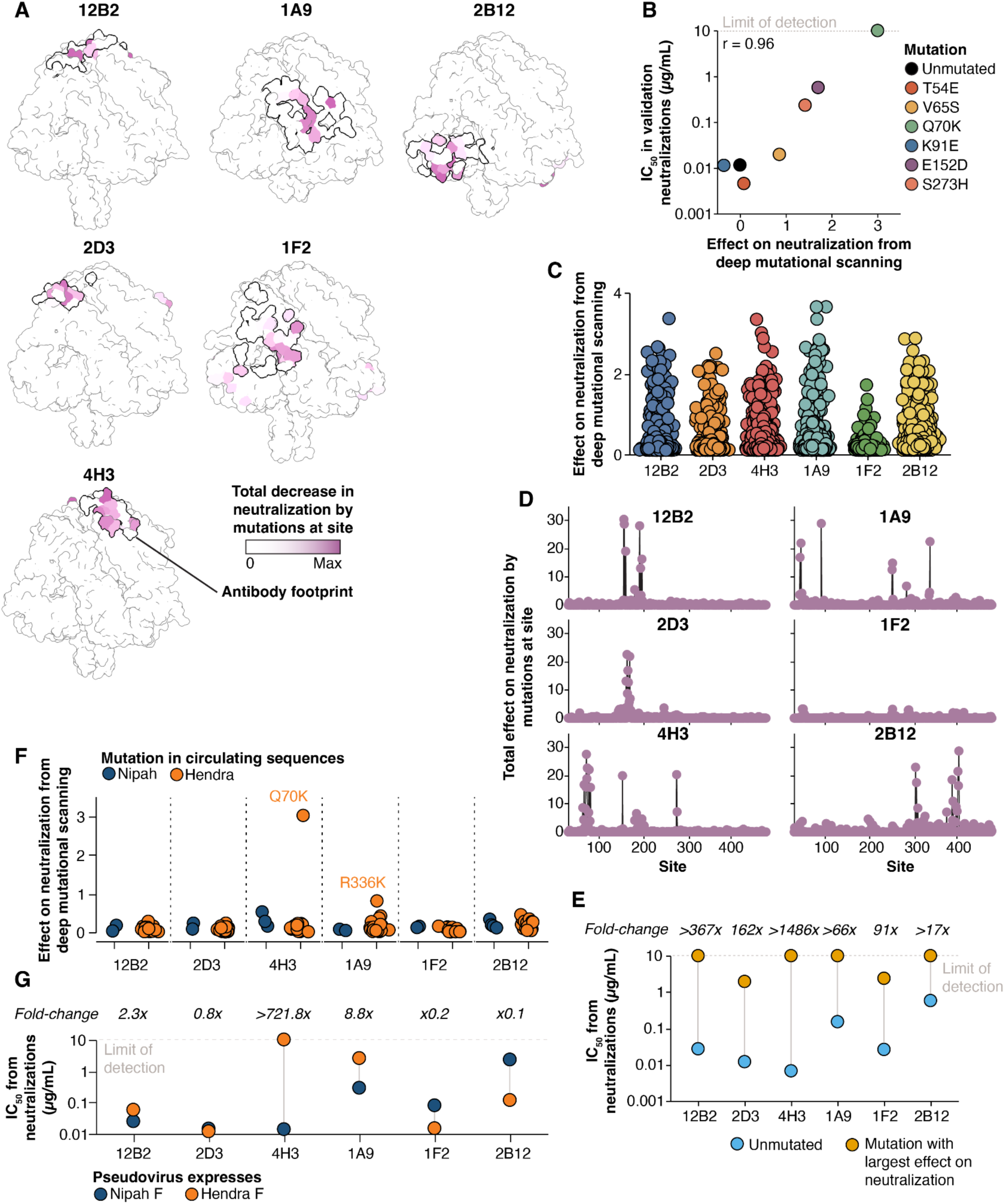
Effects of single mutations to F on antibody neutralization. **A)** Prefusion Nipah F colored by the total effect of mutations at each site on neutralization by each antibody. The thick black line shows the binding footprint of each antibody from the experimentally determined structures (see Methods for PDB accession numbers). Coloring is scaled to the maximum value separately for each antibody. See https://dms-vep.org/Nipah_Malaysia_F_DMS/visualizations/posts/AntibodyEscapeHeatmaps.html for links to interactive heatmaps that show the effects of all mutations on neutralization for each antibody. **B)** Correlation of mutation effects on neutralization by 4H3 between deep mutational scanning and traditional pseudovirus neutralization assays with the indicated F mutant. Individual mutations were selected to span a range of effects on neutralization. **C)** Extent of decreases in neutralization from each antibody caused by all mutations as measured in the deep mutational scanning. Each point is the effect of a different mutation. **D)** Total decrease in neutralization by all mutations at each site. **E)** Comparison of IC_50_ values measured in traditional neutralization assays between pseudoviruses expressing unmutated F or F with the single mutation with the greatest effect on neutralization for that antibody in the deep mutational scanning. **F)** Measured effects of all unique mutations observed in publicly available natural Nipah and Hendra F sequences on antibody neutralization in the context of Nipah F. Each point is a different amino-acid mutation observed at least twice in a Nipah or Hendra F sequence. **G)** Differences in neutralizing potency between pseudoviruses bearing either Nipah or Hendra F.

We identified mutations that affected neutralization by all antibodies, but the number and extent of the effects varied among antibodies (**Figs. 6C, 6D**). Fewer mutations affected neutralization mediated by antibody 1F2 relative to other antibodies tested: 1F2 neutralization was also affected by only a few specific amino acid changes, primarily at sites 51 and 286, while the other antibodies were affected by a much broader range of amino acid changes at multiple sites (**Figs. 6C, 6D, S10**). To confirm mutations had smaller effects on 1F2 neutralization than the other antibodies, we generated individual pseudoviruses containing the mutation identified in the deep mutational scanning data as most affecting neutralization by each antibody, and tested these pseudoviruses in traditional neutralization assays (**Figs. 6E, S11B**). While single mutations completely escaped neutralization mediated by four of the antibodies, two antibodies (1F2 and 2D3) retained detectable neutralization even against the F protein carrying the mutation that most impacted neutralization by that antibody (**Fig. 6E**).

We examined whether mutations measured in the deep mutational scanning to reduce neutralization by any of the antibodies are found in naturally occurring Nipah F sequences available in GenBank. This analysis revealed that no mutations present in circulating Nipah F sequences have strong effects on neutralization by any of the antibodies (**Fig. 6F**).

### Deep mutational scanning measurements of how mutations to Nipah F affect antibody neutralization can predict which antibodies neutralize Hendra F

To determine if our measurements of how mutations affect neutralization of Nipah virus F generalize to Hendra virus, a closely related henipavirus, we examined whether any mutations in Hendra virus F relative to Nipah virus F were measured to reduce antibody neutralization in the deep mutational scanning. One mutation found in all Hendra virus sequences (Q70K) strongly escaped antibody 4H3 neutralization in our Nipah F deep mutational scanning data, and another mutation found in all Hendra virus sequences (R336K) moderately reduced neutralization by antibody 1A9 (**Fig. 6F**). Therefore, if the deep mutational scanning measurements can be generalized from Nipah virus F to Hendra virus F, we would predict that 4H3 does not neutralize Hendra and 1A9 has modestly reduced neutralization, but the other antibodies should still neutralize Hendra virus well. To test this prediction, we generated pseudoviruses expressing Nipah virus RBP and either Nipah virus or Hendra virus F and compared their neutralization in a traditional luciferase-based assay (**Figs. 6G**, **S11C**). 4H3 did not neutralize pseudoviruses containing Hendra F even at concentrations up to 10 µg/mL, while 1A9 neutralized Hendra virus F-harboring pseudovirus with a 8.8-fold higher IC_50_ relative to Nipah virus F (**Fig. 6G**). The remaining antibodies neutralized pseudoviruses expressing either Hendra or Nipah virus F at mostly similar levels, although some antibodies had mildly reduced neutralization with Hendra virus F. To confirm that the Hendra F residues K70 and K336 were responsible for decreased sensitivity to antibodies 4H3 and 1A9, we generated pseudoviruses expressing Hendra virus F with each of these sites separately mutated to their Nipah virus F identities (K70Q or K336R) and tested their neutralization relative to pseudoviruses expressing unmutated Nipah or Hendra F. Mutating these sites in Hendra F to the Nipah virus identities greatly sensitized pseudoviruses to neutralization, resulting in IC_50_s similar to unmutated Nipah F as expected from the deep mutational scanning (**Fig. S11D)**. These results suggest our deep mutational scanning in the Nipah F background can predict which antibodies neutralize Hendra virus and identify specific mutations responsible for differences in neutralizing potency.

## Discussion

Here, we measured the effects of all single amino-acid mutations to Nipah virus F on two different protein phenotypes, cell entry and antibody neutralization. These data elucidate the constraints and evolutionary potential of this important protein, which is a key target of vaccines and therapeutics in development.

Our results show that Nipah F is less tolerant to mutations than RBP, mirroring the pattern observed in pairwise sequence identities, and more broadly with other viral entry proteins. For example, in deep mutational scanning experiments of influenza and SARS-CoV-2, which both use a single protein for binding and membrane fusion, the regions of the protein involved in fusion are generally more constrained than the regions that bind host receptors (40, 64). It appears that across diverse class I viral fusion proteins, membrane fusion is a more highly regulated and constrained phenotype than receptor binding.

Vaccine antigens based on stabilized viral entry proteins are a promising recent advance for increasing vaccine efficacy towards different viruses (60, 61, 65). Although some stabilizing mutations have been described for Nipah and Hendra virus F (29, 62, 66), we used our measurements of how mutations affect F-mediated cell entry to identify additional candidate stabilizing mutations. We focused on proline mutations in HRA due to prior work showing these mutations increase stability and do not disrupt protein folding (66). Although stabilizing mutations have been identified in other regions, such as HRB, we did not attempt to use our single-mutant deep mutational scanning data to identify candidate stabilizing mutations in HRB since stabilization in this region often requires multiple mutations (66). Other high-throughput approaches that specifically measure the effects of mutations on stability have been described for influenza virus hemagglutinin using pH as a selective agent (64, 67), since acidic pH triggers hemagglutinin-mediated fusion. Another alternative is cell-surface display approaches that directly screen for stability of the prefusion protein (68). But for our current Nipah virus F study, we were limited to using the cell entry data to identify candidate stabilizing mutations; so further experimental validation would be needed to confirm that these mutations actually increase prefusion F stability.

We also measured how all functionally tolerated F mutations affect neutralization by a panel of monoclonal antibodies, and found differences in how the neutralization by different antibodies was affected by F mutations. For instance, antibody 1F2 retained appreciable neutralization against a pseudovirus harboring F carrying the mutation that most impacted this antibody, but some other antibodies were completely escaped by single F mutations. Interestingly, some antibodies targeting HIV (69, 70) and influenza (71) have also been described that are less impacted by individual mutations. Antibodies that are more resilient to viral mutations are therefore promising candidates as Nipah virus prophylactics and therapeutics. Indeed, recent work has shown that prospective consideration of the effects of mutations during antibody development could have helped inform the development of more evolution-resistant antibody countermeasures during the COVID-19 pandemic (37).

Our study has several limitations. The experiments were performed with lentiviral pseudoviruses in cell culture. While the use of pseudoviruses obviates the substantial biosafety constraints of working with replicative Nipah virus, pseudoviruses in cell culture likely do not capture all the selective forces acting on mutations *in vivo*. Previous research on parainfluenza clinical isolates grown in cell culture demonstrated that viruses rapidly acquired substitutions in both RBP and F that altered fusogenicity (72). Our truncations to the cytoplasmic tails were necessary for effective pseudotyping, however the cytoplasmic tail of paramyxovirus RBP and F proteins have been implicated in cleavage, activation, expression and triggering (73–76). Finally, our experiments focused on measuring the effects of single amino-acid mutations to F, but in some cases the impacts of mutations during natural evolution can be modulated by epistatic interactions with other mutations (77, 78).

Overall, our study shows the Nipah F protein is more highly constrained than the RBP, making it an attractive target for vaccines and therapeutic antibodies. Our identification of highly constrained, surface-exposed residues can also help guide future attempts at defining RBP/F interacting regions, and quantifies the extent of functional constraint of different antibody epitopes. Taken together, these data shed more light on the biochemical function of F while also providing information that can help inform the development of future countermeasures to Nipah and other pathogenic henipaviruses.

## Funding and Acknowledgements

This work was supported in part by the NIH/NIAID under grant R01AI141707 to JDB, 1U19AI181881 to JDB and DV, as well as DP1AI158186, 75N93022C00036 to DV, and an Investigators in the Pathogenesis of Infectious Disease Awards from the Burroughs Wellcome Fund (DV). JDB and DV are investigators of the Howard Hughes Medical Institute and DV is the Hans Neurath Endowed Chair in Biochemistry at the University of Washington. BBL is a Washington Research Foundation Postdoctoral Fellow. SH is a postdoctoral fellow of the Translational Data Science Integrated Research Center at the Fred Hutchinson Cancer Center. This research was also supported by the Genomics & Bioinformatics Shared Resource, RRID:SCR_022606, of the Fred Hutch/University of Washington/Seattle Children’s Cancer Consortium (P30 CA015704) and the Flow Cytometry Shared Resource, RRID:SCR_022613, of the Fred Hutch/University of Washington/Seattle Children’s Cancer Consortium (P30 CA015704), and by Fred Hutch Scientific Computing, NIH grants S10-OD-020069 and S10-OD-028685. This manuscript is the result of funding in whole or in part by the National Institutes of Health (NIH). It is subject to the NIH Public Access Policy. Through acceptance of this federal funding, NIH has been given a right to make this manuscript publicly available in PubMed Central upon the Official Date of Publication, as defined by NIH.

## Author Contributions

Conceptualization: BBL, JDB

Data curation: BBL, JDB, DV

Formal analysis: BBL, DV, JDB

Funding acquisition: JDB, DV

Investigation: BBL, RG, CS

Methodology: JDB, BBL, SH

Project administration: JDB, DV

Resources: DV, JDB

Supervision: JDB, DV

Visualization: BBL, JDB

Writing - original draft: BBL, JDB

Writing - review and editing: JDB, BBL, DV

## Methods

### Data availability and interactive plots of results

All data, interactive visualizations, and raw data from experiments are publicly available on GitHub. The homepage (https://dms-vep.org/Nipah_Malaysia_F_DMS/) contains interactive visualizations to explore the deep mutational scanning data and links to additional datasets. The GitHub repository (https://github.com/dms-vep/Nipah_Malaysia_F_DMS) contains all code used to analyze the data, produce the figures, and information about available processed datasets. Sequencing data, including PacBio CCS and Illumina reads, have been deposited to the NIH SRA database under BioProject PRJNA1377637.

### Biosafety

Due to the high biosafety level and restrictions on introducing novel mutations to authentic Nipah virus, we performed all of our experiments with lentiviral-based pseudoviruses in a biosafety-level-2 laboratory by trained individuals. Importantly, these pseudoviruses are non-replicative and do not encode any other viral proteins besides Nipah F, meaning that they are not fully replicative viral pathogens. The additional proteins required for virion formation are provided *in trans* on three separate plasmids (gag/pol, tat, rev) which limits the possibility of recombination generating a replication competent virus.

We also took steps to limit information on human-specific adaptations that could be potentially misused (52, 53). Rather than perform our experiments in the context of human cells or receptors, we used specific target cells as previously described (41). Briefly, we used CHO cells that stably express bat orthologs of the Nipah receptor (ephrin-B2 or -B3) from a natural henipavirus host, *Pteropus alecto*. This approach allows us to make measurements of the antigenic and functional effects of mutations that are beneficial for countermeasure development without providing information about mutations that might potentially adapt the F protein for entry in cells expressing human receptors.

### Plasmids and primers

All plasmid and primer sequences used in this study can all be found here (https://github.com/dms-vep/Nipah_Malaysia_F_DMS/tree/main/data/paper_reference_files/sequences). All primers were ordered from Integrated DNA Technologies.

### Monoclonal antibodies

Anti-F monoclonal antibodies 4H3, 2D3, 1A9, 2B12, 1F2 were synthesized by GenScript based on the original heavy and light chain sequences (63), which are also available here (https://github.com/dms-vep/Nipah_Malaysia_F_DMS/blob/main/data/paper_reference_files/sequences/antibodies/nipahF-antibody-sequences.csv). 12B2 was produced as previously described (33).

### Cells

To produce lentiviral pseudoviruses from transfections, we used HEK-293T cells (ATCC Cat. No. CRL-3216). To produce pseudovirus from cell-stored libraries, we used HEK-293T-rtTA cells that express reverse tetracycline transactivator (rtTA) (40). HEK-293T cells were cultured in DMEM (Fisher, Cat. No. MT10013CV) supplemented with 10% Tet-free FBS (Fisher, Cat. No. A4736401), L-glutamine (Fisher Cat. No. MT25005CI), and Penicillin-Streptomycin (Fisher, Cat. No. MT30002CI).

For target cells for pseudovirus infections, we used CHO (ATCC, Cat. No. CRL-2242) cells or CHO clones expressing the bat (*Pteropus alecto*) ephrin-B2 (CHO-bEFNB2) and ephrin-B3 (CHO-bEFNB3) as previously described (41). CHO cells and bat ephrin stable CHO clones were cultured in Ham’s F-12K (Kaighn’s) Medium (Fisher, Cat. No. 21-127-030) supplemented with 10% FBS, Glutamine, and PenStrep. All cells were grown in incubators at 37°C with 5% CO2.

### Cytoplasmic tail truncations in Nipah RBP and F

To obtain accurate measurements for deep mutational scanning, high pseudovirus titers are required to limit experimental bottlenecking of library diversity. Furthermore, due to the high fusogenicity of full-length Nipah RBP and F, high levels of syncytia form between 293T producing cells during viral rescue, which can scramble the genotype-phenotype linkage in individual virions. Therefore, we generated RBP and F cytoplasmic tail truncations (**Fig. S2A**) to boost titers (**Fig. S2B**) and reduce the amount of syncytia between pseudovirus producing cells. To determine the effect of F cytoplasmic tail truncations on cleavage and activation, we performed reducing SDS-page and western blotting (See section below *Western Blotting)* between full-length and the cytoplasmic tail truncation version of F. As expected, due to the removal of the endocytosis motif in the cytoplasmic tail (76), we observed decreased amount of cleaved F (F1) in the cytoplasmic tail truncation, however there were still appreciable amounts when we rescued viruses at 48 hours after transfection (**Fig. S2C**).

### Creation of site-saturated single-mutant Nipah F libraries

We used the unmutated parental Nipah F sequence based on the reference sequence originally isolated in Malaysia during the first known outbreak in 1999 (GenBank accession NC_002728.1). We codon-optimized this sequence and removed 22 amino acids from the cytoplasmic tail at the C-terminus to improve lentiviral titers. The codon-optimized DNA sequence can be found here (https://github.com/dms-vep/Nipah_Malaysia_F_DMS/blob/main/data/paper_reference_files/sequences/unmutated_reference/NipahF_codon_optimized_CTdel_DNA.fasta).

To make the mutagenesis library we employed oligo pool (oPool) ssDNA synthesis offered by Twist BioSciences, which has an upper size limit of 300 nucleotides. The mutagenized region (sites 29-481, excluding the signal peptide and transmembrane domain) was split into six overlapping windows (**Fig. S3A**). Individual mutations at each site were designed with the most frequent human codon (For aspartic acid, we used GAT instead of GAC to limit introduction of BsmBI sites into mutant sequences) for all possible missense mutations using a custom jupyter notebook (https://github.com/dms-vep/Nipah_Malaysia_F_DMS/blob/main/notebooks/calculate_oPools.ipynb). Stop codons were designed to occur at every other position from sites 29 to 104. Each tile contained ∼1,500 specific amino-acid mutations. In order to amplify each tile and perform Golden Gate Cloning from the oPool, we also included BsmBI sites in the correct orientation on both ends of the mutagenized fragment, and unique 20bp priming sites at the extreme 3’ and 5’ ends (**Fig. S3B**). Alignments of each designed window can be found here (https://github.com/dms-vep/Nipah_Malaysia_F_DMS/tree/main/data/paper_reference_files/sequences/designed_oPools).

### PCR amplification of each tile separately from synthesized oPools

To amplify each tile separately from the delivered oPool, we used unique forward and reverse primers that matched the nucleotides added to the 3’ and 5’ ends of each tile (**Fig. S3B**). PCR primer sequences can be found here (https://github.com/dms-vep/Nipah_Malaysia_F_DMS/blob/main/data/paper_reference_files/sequences/primers/oPool_primers.csv). The PCR conditions are as follows.

25µL of 2x KOD Hot Start Master Mix (ThermoFisher, Cat. No. 71842-4), 2 µL of 10 µM of each primer, 1µL of oPool diluted DNA (0.25 ng/µL), and 20 µL of water. The thermocycler conditions were:

1. 95°C for 2 minutes
2. 95°C for 20 seconds
3. 70°C for 1 second
4. 62°C for 10s (ramp rate −0.5°C/second)
5. 68° for 25s
6. Return to step 2 for 19x cycles
7. 12°C hold.

PCR products were visualized on a 1% agarose gel to ensure each amplicon was successfully amplified and did not contain spurious bands. PCR products were cleaned with 1x AmpureXP beads (Beckman Coulter, Cat. No. A63881) and resuspended in Qiagen Buffer EB.

### Golden Gate Destination Vectors

Following PCR amplification of the individual tiles, we cloned the amplicons directly in-frame into our lentiviral vector containing unmutated Nipah F. To prepare these vectors for Golden Gate Assembly, we domesticated our lentiviral plasmid (3137_pH2rU3_ForInd_mCherry_CMV_ZsGT2APurR) by changing two pre-existing BsmBI sites by primer mutagenesis. Next, we added BsmBI sites into the Nipah F open reading frame by primer mutagenesis separately for each window, so that each mutagenized tile could be cloned directly into the vector in-frame (**Fig. S3C**). The plasmid maps of the six destination vectors can be found here (https://github.com/dms-vep/Nipah_Malaysia_F_DMS/tree/main/data/paper_reference_files/sequences/destination_vectors).

### Golden Gate Assembly of the amplified tiles and destination vectors

Each unique destination vector and amplified oPool were then combined in a Golden Gate Assembly reaction (**Fig. S3D**). The conditions are as follows. 1 µL destination vector (50fmol/µL), 1 µL PCR amplified oPool insert (100fmol/µL), 2 µL 10x T4 DNA Ligase Buffer (NEB, Cat. No. E1602S), 1 µL NEB Golden Gate Enzyme Mix BsmBI-v2 (NEB, Cat. No. E1602S), and 15 µL of water. Reactions were incubated in a thermocycler at 42°C for 1 hour, followed by 60°C for 5 minutes. Products were cleaned with 0.8x Ampure XP beads. 2 µL from each golden gate assembly reaction was electroporated into 10–beta electrocompetent cells (NEB, Cat. No. C3020K) with a BioRad MicroPulser Electroporator (Cat. No. 1652100), shocking at 2 kV, followed by resuspension of bacterial cells in 1mL of NEB 10-beta stable outgrowth media. An aliquot was diluted and plated onto LB ampicillin plates, with the remainder allowed to grow overnight at 37°C in 5mL LB and ampicillin. The total number of colony forming units (CFUs) for each unique pool ranged from 2.4e6 to 8e6, representing >1,000 CFUs per variant. High colony counts are necessary to prevent barcode swapping in subsequent steps caused by lentiviral recombination. The remaining library plasmids were extracted and purified from the overnight LB cultures with a QIAprep Spin Miniprep Kit (Qiagen, Cat. No. 27106). To ensure each mutagenized tile was inserted in-frame in the correct region, we sequenced full-length plasmids from two colonies from each reaction with Primordium.

### Barcoding the mutant libraries

After confirming the plasmids were correctly assembled, we next added a random 16 nt barcode to each plasmid, which enables us to match each Nipah F mutation with a specific barcode as previously described (**Fig. S3E**) (40, 41).

The purified plasmid libraries were pooled equimolarly based on the DNA concentrations measured by a Qubit 4 Fluorometer (ThermoFisher, Cat. No. Q33238). Plasmids were digested at 37°C for 1 hour with XbaI (NEB, Cat. No. R0145S), gel extracted with a Nucleospin Gel and PCR Clean-up kit (Macherey-Nagel, Cat. No. 740609.5), followed by purification with 0.8x Ampure XP beads. To insert unique barcodes, we setup a HiFi reaction with the XbaI digested plasmid and a ssDNA barcoding oligo (5’-gcggaactccactaggaacatttctctctcgaatctagaNNNNNNNNNNNNNNNNAGATCGGAAGAGCGTCGTGTAGGGA AAGAG-3’). The HiFi conditions are as follows. 5 µL Barcoding oligo (0.2 pmol), 1 µL digested vector (10 fmol), 4 µL of water, and 10 µL of 2x HiFi MasterMix (NEB, Cat. No. E2621L). Reactions were incubated at 50°C for 1 hour, followed by electroporation into NEB 10-beta electrocompetent cells. To create separate replicate libraries, we used two separate HiFi reactions which became LibA and LibB. Electroporations were grown in LB and ampicillin overnight, with a small aliquot diluted to count the number of transformants. We obtained a total of 3.6e6 CFU for LibA, and 6.2e6 for LibB, which is >400 CFU per barcoded variant. As mentioned previously, the number of colonies needs to be significantly higher than the total size of the library to ensure enough unique barcodes are present to avoid swapping during lentiviral recombination. After overnight growth, plasmids were extracted and purified with a Qiagen Spin MiniPrep kit.

### Producing genotype-phenotype linked mutant pseudoviruses

We produced genotype-phenotype linked pseudoviruses as previously described(40, 41). Our method relies on coupling the genotype contained within the lentiviral genome with the mutagenized proteins on the surface of the pseudovirus. To accomplish this, we rescued VSV-G pseudotyped viruses containing the mutagenized lentiviral genome, followed by infection at low MOI to ensure each cell has no more than a single integrated provirus. This ensures that pseudoviruses produced from each individual cell are all genotype-phenotyped linked. Details of our method are as follows: we transfected HEK293T with the Nipah F library cloned into a lentiviral backbone (described above), plus three lentiviral helper plasmids (26_HDM_Hgpm227, 27_HDM_tat1b, 28_pRC_CMV_Rev1b), an expression plasmid for VSV-G (29_HDM_VSV_G), and BioT transfection reagent (Biolands Scientific, Cat. No. B01-02). Plasmid sequences can be found here: (https://github.com/dms-vep/Nipah_Malaysia_F_DMS/tree/main/data/paper_reference_files/sequences/plasmids). Supernatants were filtered through a 0.45 µm syringe filter (Corning, Cat. No. 431220) after 48 hours and titered on HEK293T-rtTA(40) cells to estimate an appropriate MOI. HEK293T-rtTA cells were then infected with pseudoviruses expressing VSV-G at an MOI of <0.01 to ensure each cell contained no more than a single integrated lentivirus. Cells were then passaged in the presence of 1 µg/mL puromycin (ThermoFisher Cat. No. A1113803) until all cells were ZsGreen positive. These cell-stored libraries were frozen down with 10% DMSO and stored in the gas phase of liquid nitrogen.

To generate the library virus used for infections, we thawed a fresh aliquot of cells and cultured them until they could be plated onto multiple five-layer flasks (Corning Falcon 875cm2 Rectangular Neck Cell Culture Multi-Flask, Cat. No. 353144). The following day, we transfected the cells. To generate VSV-G expressing pseudoviruses from the library, we prepared the following transfection mix. 45 µg each of the three helper plasmids (gag/pol, tat, rev), 15 µg of VSV-G expression plasmid, 7.5 mL of DMEM, and 225 µL of BioT transfection reagent. Transfection mixtures were incubated for 10 minutes at room temperature and added to each five-layer flask. 48 hours later, supernatants were filtered through a 0.45 µm SFCA Nalgene 500 mL Rapid-Flow filter unit (Cat. No. 09-740-44B).

To produce the RBP_wt_/F_mutant_ pseudovirus libraries, the same approach above was used, except we used 48.75 µg of each helper plasmid, and 3.75 µg of unmutated Nipah RBP (3336_HDM_Nipah_RBP_CTdel, RBP_wt_) that was codon-optimized and contained a 32 amino-acid truncation in the N-terminal cytoplasmic tail(41). Titers of unconcentrated library pseudovirus were typically around 5*10^4^ transducing units (TU)/mL. Following filtration of the RBP_wt_/F_mutant_ pseudovirus libraries, we concentrated them to obtain high titers needed for our selections.

We used two different methods to concentrate the lentiviruses for selections, which did not differ in their concentration efficiency or affect the results. We mixed virus supernatant and Lenti-X Concentrator (Takara, Cat. No. 631232) at a 3:1 ratio and incubated overnight at 4°C. Tubes were then spun at 1500g for 45 minutes at 4°C. Supernatants were discarded, and pellets were resuspended in complete F-12K media. Alternatively, we added 50mL of virus supernatants to Pierce Protein Concentrators, 100k MWCO (Thermo, Cat. No. 88537) and spun at 2000g for 30 minutes. Liquid at the top of the filters containing the concentrated pseudoviruses was transferred to new tubes. All concentrated viruses were stored in aliquots at −80°C. Following concentration, titers were typically around 1*10^6^ TU/mL.

### Library infections and DNA template recovery

To obtain DNA templates from lentiviruses for PCR amplification and sequencing, we used the following approach as previously described (40, 41). Briefly, our method relies on obtaining unintegrated lentiviral DNA that is generated by lentiviral reverse transcription. This serves two main purposes. First, most HIV DNA templates do not integrate into the host genome following infection. Second, if we attempted to extract provirus from cell gDNA, it would decrease PCR efficiency. Therefore, we infect target cells with 1-2×10^6^ TU of library virus and 8-12 hours later extract low molecular weight DNA using a Qiagen QIAamp Spin Miniprep kit. For VSV-G pseudotyped lentiviruses, we infected CHO cells plated in six-well dishes, for RBP_wt_/F_mutant_ pseudoviruses we either used CHO-bEFNB2 or -bEFNB3 cells. 8-12 hours later, media was removed from the cells, and 450µL of trypsin was added. After 5 minutes, 550µL of complete F-12K media was added and mixed to dislodge all the cells. Cells were pelleted at 300g for 4 minutes. Supernatant was removed and cells were resuspended in 250µL of P1 Buffer. Cell suspensions were then miniprepped to isolate non-integrated lentiviral templates.

### Long-read sequencing to link mutations to barcodes

To link the 16 nt random barcodes with each mutation, we performed long-read PacBio sequencing of lentiviral DNA templates that were miniprepped from target cells 8-12 hours after infection with the concentrated VSV-G expressing library virus as previously described (40, 41). Due to the pseudodiploid nature of lentiviruses and possible recombination, the linkage of barcodes and mutations is performed after generating the cell libraries to ensure the linkage is accurate. Our approach here includes low PCR cycles to minimize strand exchange, while also quantifying the amount of strand exchange during PCR, which artificially scrambles the barcode-mutation linkages. Specifically, we included single nucleotide tags (either ‘G’ or ‘C’) within the first round primers. The amplicons from the first round PCRs are then pooled and undergo a second round of PCR. Strand exchange that occurred during the 2nd round of PCR is identified by comparing the single nucleotide tags at the 5’ and 3’ ends, which were introduced in the first round PCR.

1st round PCRs were done in three separate reactions to add either a G or C at the 5’ and 3’ ends to allow identification of strand exchange. 1st round PCR conditions were as follows. 20 µL of KOD Hot Start Master Mix, 10µL of DNA templates from minipreps, 1µL each of 10µM primers (PacBio_5pri_G + PacBio_3pri_C) or (PacBio_5pri_C + PacBio_3pri_G), and 8µL of water. The thermocycler conditions were:

1. 95°C for 2 minutes
2. 95°C for 20 seconds
3. 70°C for 1 second
4. 60°C for 10 seconds with a ramp rate of −0.5°C/sec
5. 70°C for 1 minute
6. Return to step 2 for 7x cycles
7. 12°C hold

PCR products from three separate first round PCR were pooled and cleaned up with 1x Ampure XP beads. The conditions for the 2nd round PCR are as follows. 25 µL of KOD Hot Start Master Mix, 2 µL of each primer at 10 µM (5’_PB_Rnd2 and 3’_PB_Rnd2), and 21 µL of cleaned round 1 product. The thermocycler conditions were:

1. 95°C for 2 min
2. 95°C for 20 seconds
3. 70°C for 1 second
4. 60°C for 10 seconds with a ramp rate of −0.5°C/sec,
5. 70°C for 1 minute.
6. Return to step 2 for 10x cycles
7. 12°C hold

PCR products were then cleaned with 1x Ampure XP beads, followed by circular consensus sequencing on a PacBio Sequel IIe. We found that 2-3% of all reads had strand exchange, as identified by comparing the single nucleotide tags described above.

### Illumina barcode sequencing

To recover barcode frequencies associated with each variant, we performed Illumina sequencing on amplified unintegrated viral template DNA extracted from target cells 8-12 hours after infection (see section *Library infections and DNA template recovery*). Next, we set up PCRs for each selection experiment with the following conditions. 25 µL of KOD Hot Start Master Mix, 1.5 µL each of forward and reverse 1st round Illumina primers (Illumina_Rnd1_For and Illumina_Rnd1_Rev3), and 22 µL of the miniprepped DNA templates. The thermocycler conditions were:

1. 95°C for 2 minutes
2. 95°C for 20 seconds
3. 70°C for 1 second
4. 58°C for 10 seconds with a ramp rate of −0.5°C/sec
5. 70°C for 20 seconds
6. Return to step 2 for 27x cycles
7. 12°C hold

Following 1st round PCRs, we measured DNA concentrations using a Qubit 4 Fluorometer. DNA concentrations were normalized and then added to 2nd round PCRs each containing 2 µL of 10 µM unique indexed 2nd-round primers, which is used to identify index-hopping that can occur during Illumina sequencing(79). PCR conditions were similar to above, except only 1 µL of 1st round PCR was used as template, with the remaining volume containing water, and only 20 PCR cycles were used for the 2nd round PCRs. Next, the indexed 2nd round samples were pooled equimolarly and run out on a 1% agarose gel. DNA from bands of the correct size were excised from the gels and extracted with a Nucleospin Gel and PCR Clean-up kit. The DNA was then cleaned with a final purification step using 0.8x Ampure XP beads and washed three times with 80% EtOH. Pooled samples were then sequenced on an Illumina NextSeq with a P2 kit, or a NovaSeq lane for 50 cycles, depending on how many multiplexed, indexed samples we included. We typically received 20-100 million reads per sample, corresponding to >500x coverage of each variant.

### Analysis of PacBio sequencing and creation of barcode-variant lookup table

For processing the sequencing data from the deep mutational scanning experiments, we used the *dms-vep-pipeline-3 v3.25.0* package (https://github.com/dms-vep/dms-vep-pipeline-3), which is briefly described here. To link specific mutations with each barcode, we performed PacBio circular consensus sequencing (CCS) on amplicons spanning the entire F protein and the 16 nt barcodes (see section *PacBio sequencing*).

From the PacBio CCS data obtained for both libraries, we first aligned the reads to the unmutated Nipah F reference sequence using the *alignparse* package (80). Reads that aligned poorly or had a higher than expected number of mutations in the unmutated regions were filtered out. Next, variants that did not contain a barcode or were the result of strand exchange were filtered out. Consensus sequences for each barcode/variant sequence were constructed using *alignparse*, while requiring a minimum of at least three CCS reads and a max cutoff of 0.2 for any minor variants within the consensus. The final barcode/variant lookup tables were used as a reference for all downstream analyses that used the short-read Illumina sequencing of the barcodes only.

### Calculation of functional scores

To calculate the efficiency of cell entry of different variants relative to the unmutated sequence, we compared variant frequencies derived from infecting cells with either VSV-G or RBP_wt_/F_mutant_ pseudotyped viruses as previously described(40). Here, the VSV-G condition serves as a ‘control’, and allows us to determine frequencies of variants contained within the lentiviral genome that would be highly deleterious for F-mediated entry. From Illumina sequencing of barcodes recovered from these infections, we obtain relative frequencies of each infecting variant.

Illumina sequencing data were first filtered to ensure that all bases had a minimum sequencing quality score of 20, and were then aligned to the barcode variant table generated from the PacBio CCS described above. Barcode sequences below a frequency 2*10^-5^ were excluded from downstream analyses.

We next compared the frequency of barcodes between the VSV-G and RBP_wt_/F_mutant_ selections using the package *dms_variants* (https://github.com/jbloomlab/dms_variants) as previously described (40, 41). Briefly, functional scores were calculated using enrichment ratios: log_2_([n^v^_post_ / n^wt^_post_] / [n^v^_pre_/n^wt^_pre_]) where n^v^_post_ and n^wt^_post_ are the counts of variant *v* or the unmutated variants from the Nipah F pseudovirus infection, respectively. The variant *v* or unmutated counts in the VSV-G pseudotyped infection are n^v^_pre_ and n^wt^_pre_, respectively. The lower limit of detection for mutation effects was approximately −4, which corresponds to the median effects of stop codons, which should result in non-functional protein. Therefore, we clipped the functional scores below this threshold and used those functional scores for our cell entry calculations.

### Calculation of mutation effects on cell entry

Although our libraries primarily contain variants with a single mutation relative to the unmutated parental strain (69%), a subset of our library contains multiple amino-acid mutations. To decompose the effects of these multi-mutants, we utilized the *multiDMS* package to apply global epistasis fitting (49, 81). We compared the effects of the decomposed effects with the uncorrected functional scores of single-mutations only (**Fig. S5**). These comparisons were all well-correlated, with outlier mutations generally having few (<2) barcodes associated with a single mutation. Thus, for all downstream analyses, we used the decomposed functional effects produced from the global epistasis modeling.

To generate the final cell entry effect values for each mutation, we performed a total of two technical replicates from two separate rescues for each library, for a total of eight functional selections (four each from LibA and LibB). The effects for each mutation were then averaged from the eight functional selections. To filter out low quality or noisy data, we applied two filters. First, we required each mutation to occur with two unique barcodes (*times_seen >= 2*). Second, we removed any mutation that had a high standard deviation between replicates (*effect_std <= 1*). The final functional effect values reported in the figures correspond to the average effect across libraries and replicates.

### Effects of mutations on antibody neutralization

To measure the effects of mutations on antibody neutralization, we used a previously described method (40) with a few modifications described here. ∼1×10^6^ CHO-bEFNB3 cells were plated in individual wells of a 6-well plate. The following day, ∼1×10^6^ TU of library virus was either added directly onto the cells (no-antibody control) or were incubated in the presence of antibody for one hour prior to adding to cells. Antibody concentrations were selected that generally corresponded to a range at which 50% of variants were neutralized, up to 99.5%. During DNA template extraction, we spiked in DNA plasmid containing eight known barcodes that would correspond to ∼1% of the reads in the no-antibody control. This DNA plasmid spike-in allowed us to estimate the amount of neutralization each antibody condition had relative to the no-antibody control, as previously described (40).

Following extraction and sequencing of the barcodes, we parsed, filtered, and aligned the barcodes as described in the previous section. We calculated an escape score as the log_2_ transformed values of: (F * [n^v^_post_ * N_pre_] / [n^v^_pre_ * N_post_]) where F is the overall fraction of the library that escapes neutralization, which is derived from the known DNA barcodes that we spiked in during template extraction. n^v^_post_ and n^v^_post_ are the counts of variant *v* in the antibody condition and no-antibody condition, respectively. N_pre_ and N_post_ are the summed counts of all variants in the no-antibody control and antibody condition, respectively. We then fit neutralization curves for each selection as implemented in the package *polyclonal* (https://jbloomlab.github.io/polyclonal/) (82). We filtered the antibody escape data using three different cutoffs. First, we required mutations to occur with two unique barcodes (*times_seen >=2).* Second, since we can only measure mutations that have at least some level of cell entry, we excluded measurements of mutations with very low cell entry scores (*min_func_effect >= −2.5).* Third, we filtered out mutations that had high standard deviations between replicates (*escape_std_dev <= 2).* Reported escape scores for each mutation are the average effect calculated from at least two different independent selections with LibA and LibB.

### Validation of mutation effects using individual pseudoviruses

To validate the effects of mutations on cell-entry and antibody neutralization, we generated a set of plasmids expressing Nipah F that contained different single mutations. The parental Nipah F sequence is identical to the unmutated Nipah F sequence used in the lentiviral vector described above, but instead placed into a mammalian expression vector (5073_HDM_NipahF_CTdel_GeneArt). All single amino-acid mutations were generated by primer mutagenesis followed by sequencing confirmation by Primordium.

For cell entry validations, we selected mutations spanning a range of entry effects, which include K98R, L104N, V108D, R109Q, G215R, and E361G, which corresponded to a range of predicted cell entry effects. We made three separate plasmid preps of each Nipah F validation mutation and transfected them into 293T cells, along with plasmid 2727_pHAGE6_Luciferase, 26_HDM_Hgpm2, and 3336_HDM_Nipah_RBP_CTdel. 48 hours later, supernatants were filtered to remove cell debris. CHO-bEFNB3 cells plated on a 96-well plate were infected with dilutions of virus supernatant for each mutation and replicate, and Luciferase measurements were taken 48 hours later on a plate reader using a Bright-Glo Luciferase Assay System (Promega, Cat. No. E2620). We then checked to make sure our readings were within a linear range and compared the average luciferase signal of each mutation to the unmutated Nipah F readings.

To validate the magnitude of escape among antibodies, we generated the top escape mutation that had a cell entry score greater than −1 for each antibody in the same mammalian expression vector described above. These mutations were V159R (12B2), A165L (2D3), V65D (4H3), T43P (1A9), T286W (1F2), and E406T (2B12). To validate 4H3 escape measurements, we made mutations corresponding to Nipah F T54E, K98R, T250I, T250W, D255E, F282T, and N350E. We used a luciferase-based system identical to the cell entry assay described above with the following differences. Filtered virus supernatants were incubated with different concentrations of antibody for one hour prior to adding to 96-well plates containing CHO-bEFNB3 cells. Luciferase measurements were taken identical to above. Relative luciferase readings compared to wells without antibody were used to generate neutralization curves. Neutralization assays were performed in duplicate, unless noted otherwise.

### Western Blotting

To determine the effects of the cytoplasmic tails truncations on F cleavage and RBP/F incorporation into pseudoviruses, we performed reducing SDS-PAGE followed by western blotting. To generate the pseudovirus, we transfected 15cm plates containing 293T cells with 15 µg of plasmid 2727_pHAGE6_Luciferase, 10µg of plasmid 26_HDM_Hgpm2, 45 µL BioT, and combinations of RBP+F plasmids (2.5µg). 30 hours later, supernatants were filtered through a 0.45 µm syringe filter (Corning, Cat. No. 431220), followed by ulltracentrifugation at 100,000 x g for 1 hour. Pellets were resuspended in 500 µL of PBS. An aliquot of the concentrated virus was mixed with 5x Pierce Lane Marker Reducing Sample (ThermoFisher Cat. No. 39000) and boiled for 3 minutes. 10 µL of protein or 5 µL of marker (BioRad Precision Plus Protein Dual Color Standards; Cat. No. 1610374) were loaded onto SDS-PAGE gels (4-20% Mini-PROTEAN TGX Precast Protein Gels, 10-well, 50 µL; BioRad Cat. No. 4561094) and run at 100V for 1.5 hours. Gels were transferred with iBlot3 Transfer Stacks Mini PVDF (Invitrogen Cat. No. IB34002). Membranes were blocked for 2 hours with 5% milk in 1X Tris Buffered Saline with Tween 20. Primary antibodies were then used to stain for p24 at 1:4000 (Rabbit anti-HIV1 p24 antibody; Abcam Cat. No. ab32352), RBP at 1:3000 (Rabbit pAb Nipah virus Glycoprotein G; SinoBiological Cat. No. 40980-T62), or F at 1:3000 (Rabbit Anti-Nipah Fusion F0 polyclonal antibody; Antibody system Cat. No. PVV08101). After 2 hours, membranes were washed 3x with TBST, followed by incubation with a goat anti-rabbit HRP secondary antibody (Invitrogen Cat. No. 31460) at 1:5000. After 1 hour of gentle shaking, membranes were washed 3x with TBST and 2X with PBS, followed by developing with SuperSignal West Pico PLUS Chemiluminescent Substrate (ThermoScientific Cat. No. 34580). After 5 minutes, membranes were imaged and analyzed in ImageJ.

### Sequence analysis of publicly available sequences

To generate sequence alignments, we downloaded all complete whole genomes from Nipah and Hendra viruses from GenBank (accessed May 6th, 2025). GenBank accessions of all sequences can be found here (https://github.com/dms-vep/Nipah_Malaysia_F_DMS/tree/main/data/paper_reference_files/sequences/genbank). Sequences were aligned using MAFFT v7.520 (83), and maximum-likelihood phylogenetic trees were inferred with IQ-Tree v2.2.2.6 (84). Alignments corresponding to RBP and F were extracted using Geneious Prime v2024.0.7. Amino-acid polymorphisms were calculated from the alignments using a custom jupyter notebook (https://github.com/dms-vep/Nipah_Malaysia_F_DMS/blob/main/analysis/workflow/notebooks/find_variable_sites_from_alignments.ipynb). Phylogenetic trees were visualized with the *baltic v0.3* package (https://github.com/evogytis/baltic).

### Structural analyses

All protein structures were visualized with ChimeraX v1.10 (85) using publicly available structures deposited in the Protein Data Bank (PDB). For the Nipah F prefusion structure, we used PDB 5EVM. For the antibody structures, we used PDBs 7UPK (1A9), 7UPD (2B12), 7UP9 (2D3), 7UOP (4H3), and 7KI4 (12B2). For 1F2, there is not a structure available, but based on previous low-resolution cryo-EM, the binding interface is very similar to antibody 1H8 (7UPA) (63), which was used for 1F2 footprint estimates. The Nipah virus F postfusion structure was generated with AlphaFold2 (86) through CollabFold (87) using the postfusion structure of Langya virus F (8TVE) (21) as a template before removing residues with the lowest confidence from the model manually.

### Identification of candidate prefusion stabilizing mutations

To identify candidate prefusion stabilizing mutations, we reasoned that sites with highly deleterious proline mutations in regions that undergo large conformational changes between the pre- and postfusion conformations restrict the transition to the postfusion conformation, which is necessary for cell entry. However, if all or most other mutations at a site are deleterious, the unmutated residue is likely critical for correct protein folding and is not a good candidate for introducing stabilizing mutations. We used the cell entry deep mutational scanning data to identify sites meeting specific filtering criteria, with the rationale for each given below. We only included sites within the apical domain of prefusion F, since it undergoes the largest change between pre- and postfusion conformations (59), and previously described stabilizing mutations were all located in the apical domain (29, 62). We excluded sites in the fusion peptide (sites 110-125) since they are likely important for membrane fusion. Additional criteria included the site could not be in a helix or sheet in the prefusion structure, or at a residue that is involved in hydrogen bonding with sidechains in nearby residues, as these sites are likely important for maintaining correct protein folding. We required the proline mutation to have a cell entry score <-3, and the site needed at least four mutations with a cell entry score >-1. Finally, the site could not be at a N-linked glycosylation site or cause a disruption in a glycosylation site (i.e. a proline mutation at X in the NX(T/S) glycosylation motif).

## References

1. K. B. Chua, et al., Nipah virus: a recently emergent deadly paramyxovirus. Science 288, 1432–1435 (2000).

2. S. P. Luby, et al., Recurrent zoonotic transmission of Nipah virus into humans, Bangladesh, 2001-2007. Emerg. Infect. Dis. 15, 1229–1235 (2009).

3. J. H. Epstein, H. E. Field, S. Luby, J. R. C. Pulliam, P. Daszak, Nipah virus: impact, origins, and causes of emergence. Curr. Infect. Dis. Rep. 8, 59–65 (2006).

4. K. B. Chua, et al., Fatal encephalitis due to Nipah virus among pig-farmers in Malaysia. Lancet 354, 1257–1259 (1999).

5. Centers for Disease Control and Prevention (CDC), Outbreak of Hendra-like virus--Malaysia and Singapore, 1998-1999. MMWR Morb. Mortal. Wkly. Rep. 48, 265–269 (1999).

6. B. Nikolay, et al., Transmission of Nipah Virus - 14 Years of Investigations in Bangladesh. N. Engl. J. Med. 380, 1804–1814 (2019).

7. E. S. Gurley, et al., Person-to-person transmission of Nipah virus in a Bangladeshi community. Emerg. Infect. Dis. 13, 1031–1037 (2007).

8. C. T. Pager, W. W. Craft Jr, J. Patch, R. E. Dutch, A mature and fusogenic form of the Nipah virus fusion protein requires proteolytic processing by cathepsin L. Virology 346, 251–257 (2006).

9. S. Diederich, et al., Activation of the Nipah virus fusion protein in MDCK cells is mediated by cathepsin B within the endosome-recycling compartment. J. Virol. 86, 3736–3745 (2012).

10. S. Diederich, E. Dietzel, A. Maisner, Nipah virus fusion protein: influence of cleavage site mutations on the cleavability by cathepsin L, trypsin and furin. Virus Res. 145, 300–306 (2009).

11. S. Diederich, L. Thiel, A. Maisner, Role of endocytosis and cathepsin-mediated activation in Nipah virus entry. Virology 375, 391–400 (2008).

12. O. Pernet, Y. E. Wang, B. Lee, Henipavirus receptor usage and tropism. Curr. Top. Microbiol. Immunol. 359, 59–78 (2012).

13. T. A. Bowden, et al., Structural basis of Nipah and Hendra virus attachment to their cell-surface receptor ephrin-B2. Nat. Struct. Mol. Biol. 15, 567–572 (2008).

14. K. Xu, et al., Host cell recognition by the henipaviruses: crystal structures of the Nipah G attachment glycoprotein and its complex with ephrin-B3. Proc. Natl. Acad. Sci. U. S. A. 105, 9953–9958 (2008).

15. O. A. Negrete, et al., EphrinB2 is the entry receptor for Nipah virus, an emergent deadly paramyxovirus. Nature 436, 401–405 (2005).

16. R. A. Lamb, R. G. Paterson, T. S. Jardetzky, Paramyxovirus membrane fusion: lessons from the F and HN atomic structures. Virology 344, 30–37 (2006).

17. A. Chang, R. E. Dutch, Paramyxovirus fusion and entry: multiple paths to a common end. Viruses 4, 613–636 (2012).

18. C. J. Russell, L. E. Luque, The structural basis of paramyxovirus invasion. Trends Microbiol. 14, 243–246 (2006).

19. D. Kalbermatter, et al., Structure and supramolecular organization of the canine distemper virus attachment glycoprotein. Proc. Natl. Acad. Sci. U. S. A. 120, e2208866120 (2023).

20. Z. Wang, et al., Architecture and antigenicity of the Nipah virus attachment glycoprotein. Science 375, 1373–1378 (2022).

21. Z. Wang, et al., Structure and design of Langya virus glycoprotein antigens. Proc. Natl. Acad. Sci. U. S. A. 121, e2314990121 (2024).

22. B. D. Welch, et al., Structure of the parainfluenza virus 5 (PIV5) hemagglutinin-neuraminidase (HN) ectodomain. PLoS Pathog. 9, e1003534 (2013).

23. P. Yuan, et al., Structure of the Newcastle disease virus hemagglutinin-neuraminidase (HN) ectodomain reveals a four-helix bundle stalk. Proc. Natl. Acad. Sci. U. S. A. 108, 14920–14925 (2011).

24. J. J. W. Wong, R. G. Paterson, R. A. Lamb, T. S. Jardetzky, Structure and stabilization of the Hendra virus F glycoprotein in its prefusion form. Proc. Natl. Acad. Sci. U. S. A. 113, 1056–1061 (2016).

25. K. Xu, et al., Crystal Structure of the Pre-fusion Nipah Virus Fusion Glycoprotein Reveals a Novel Hexamer-of-Trimers Assembly. PLoS Pathog. 11, e1005322 (2015).

26. T. C. Marcink, et al., Subnanometer structure of an enveloped virus fusion complex on viral surface reveals new entry mechanisms. Sci Adv 9, eade2727 (2023).

27. E. C. Smith, A. Popa, A. Chang, C. Masante, R. E. Dutch, Viral entry mechanisms: the increasing diversity of paramyxovirus entry: The increasing diversity of paramyxovirus entry. FEBS J. 276, 7217–7227 (2009).

28. R. M. Iorio, P. J. Mahon, Paramyxoviruses: different receptors -different mechanisms of fusion. Trends Microbiol. 16, 135–137 (2008).

29. R. J. Loomis, et al., Structure-Based Design of Nipah Virus Vaccines: A Generalizable Approach to Paramyxovirus Immunogen Development. Front. Immunol. 11, 842 (2020).

30. Y.-P. Chan, et al., Biochemical, conformational, and immunogenic analysis of soluble trimeric forms of henipavirus fusion glycoproteins. J. Virol. 86, 11457–11471 (2012).

31. L. Zeitlin, et al., Therapeutic administration of a cross-reactive mAb targeting the fusion glycoprotein of Nipah virus protects nonhuman primates. Sci. Transl. Med. 16, eadl2055 (2024).

32. C. E. Mire, et al., A cross-reactive humanized monoclonal antibody targeting fusion glycoprotein function protects ferrets against lethal nipah virus and Hendra virus infection. J. Infect. Dis. 221, S471–S479 (2020).

33. H. V. Dang, et al., Broadly neutralizing antibody cocktails targeting Nipah virus and Hendra virus fusion glycoproteins. Nat. Struct. Mol. Biol. 28, 426–434 (2021).

34. H. V. Dang, et al., An antibody against the F glycoprotein inhibits Nipah and Hendra virus infections. Nat. Struct. Mol. Biol. 26, 980–987 (2019).

35. A. J. Greaney, et al., Complete Mapping of Mutations to the SARS-CoV-2 Spike Receptor-Binding Domain that Escape Antibody Recognition. Cell Host Microbe 29, 44–57.e9 (2021).

36. A. Moulana, et al., The landscape of antibody binding affinity in SARS-CoV-2 Omicron BA.1 evolution. Elife 12 (2023).

37. F. Jian, et al., Viral evolution prediction identifies broadly neutralizing antibodies to existing and prospective SARS-CoV-2 variants. Nat. Microbiol. 10, 2003–2017 (2025).

38. J. L. R. Zamora, et al., Third Helical Domain of the Nipah Virus Fusion Glycoprotein Modulates both Early and Late Steps in the Membrane Fusion Cascade. J. Virol. 94 (2020).

39. R. K. Plemper, R. W. Compans, Mutations in the putative HR-C region of the measles virus F2 glycoprotein modulate syncytium formation. J. Virol. 77, 4181–4190 (2003).

40. B. Dadonaite, et al., A pseudovirus system enables deep mutational scanning of the full SARS-CoV-2 spike. Cell 186, 1263–1278.e20 (2023).

41. B. B. Larsen, et al., Functional and antigenic landscape of the Nipah virus receptor-binding protein. Cell 188, 2480–2494.e22 (2025).

42. D. Khetawat, C. C. Broder, A functional henipavirus envelope glycoprotein pseudotyped lentivirus assay system. Virol. J. 7, 312 (2010).

43. S. R. Witting, P. Vallanda, A. L. Gamble, Characterization of a third generation lentiviral vector pseudotyped with Nipah virus envelope proteins for endothelial cell transduction. Gene Ther. 20, 997–1005 (2013).

44. C. B. Macdonald, et al., DIMPLE: deep insertion, deletion, and missense mutation libraries for exploring protein variation in evolution, disease, and biology. Genome Biol. 24, 36 (2023).

45. N. Daffern, I. M. Francino-Urdaniz, Z. T. Baumer, T. A. Whitehead, Standardizing cassette-based deep mutagenesis by Golden Gate assembly. Biotechnol. Bioeng. 121, 281–290 (2024).

46. B. Álvarez-Rodríguez, S. Velandia-Álvarez, C. Toft, R. Geller, Mapping mutational fitness effects across the coxsackievirus B3 proteome reveals distinct profiles of mutation tolerability. PLoS Biol. 22, e3002709 (2024).

47. J. M. O. Rawson, O. A. Nikolaitchik, B. F. Keele, V. K. Pathak, W.-S. Hu, Recombination is required for efficient HIV-1 replication and the maintenance of viral genome integrity. Nucleic Acids Res. 46, 10535–10545 (2018).

48. A. E. Jetzt, et al., High rate of recombination throughout the human immunodeficiency virus type 1 genome. J. Virol. 74, 1234–1240 (2000).

49. J. Otwinowski, D. M. McCandlish, J. B. Plotkin, Inferring the shape of global epistasis. Proc. Natl. Acad. Sci. U. S. A. 115, E7550–E7558 (2018).

50. M. I. Bonaparte, et al., Ephrin-B2 ligand is a functional receptor for Hendra virus and Nipah virus. Proc. Natl. Acad. Sci. U. S. A. 102, 10652–10657 (2005).

51. O. A. Negrete, et al., Two key residues in ephrinB3 are critical for its use as an alternative receptor for Nipah virus. PLoS Pathog. 2, e7 (2006).

52. K. M. Esvelt, Inoculating science against potential pandemics and information hazards. PLoS Pathog. 14, e1007286 (2018).

53. G. Lewis, P. Millett, A. Sandberg, A. Snyder-Beattie, G. Gronvall, Information Hazards in Biotechnology. Risk Anal. 39, 975–981 (2019).

54. H. C. Aguilar, V. Aspericueta, L. R. Robinson, K. E. Aanensen, B. Lee, A quantitative and kinetic fusion protein-triggering assay can discern distinct steps in the nipah virus membrane fusion cascade. J. Virol. 84, 8033–8041 (2010).

55. C. J. Russell, T. S. Jardetzky, R. A. Lamb, Conserved glycine residues in the fusion peptide of the paramyxovirus fusion protein regulate activation of the native state. J. Virol. 78, 13727–13742 (2004).

56. E. C. Smith, S. M. Gregory, L. K. Tamm, T. P. Creamer, R. E. Dutch, Role of sequence and structure of the Hendra fusion protein fusion peptide in membrane fusion. J. Biol. Chem. 287, 30035–30048 (2012).

57. M. Moll, S. Diederich, H.-D. Klenk, M. Czub, A. Maisner, Ubiquitous activation of the Nipah virus fusion protein does not require a basic amino acid at the cleavage site. J. Virol. 78, 9705–9712 (2004).

58. Q. Wang, et al., The nanoscale organization of the Nipah virus fusion protein informs new membrane fusion mechanisms. Elife 13 (2025).

59. D. S. Zyla, et al., A neutralizing antibody prevents postfusion transition of measles virus fusion protein. Science 384, eadm8693 (2024).

60. J. E. Bowen, et al., SARS-CoV-2 spike conformation determines plasma neutralizing activity elicited by a wide panel of human vaccines. Sci. Immunol. 7, eadf1421 (2022).

61. M. C. Crank, et al., A proof of concept for structure-based vaccine design targeting RSV in humans. Science 365, 505–509 (2019).

62. P. O. Byrne, et al., Prefusion stabilization of the Hendra and Langya virus F proteins. J. Virol. 98, e0137223 (2024).

63. P. O. Byrne, et al., Structural basis for antibody recognition of vulnerable epitopes on Nipah virus F protein. Nat. Commun. 14, 1494 (2023).

64. T. C. Yu, et al., Pleiotropic mutational effects on function and stability constrain the antigenic evolution of influenza hemagglutinin. bioRxivorg 2025.05.24.655919 (2025).

65. J. S. McLellan, et al., Structure-based design of a fusion glycoprotein vaccine for respiratory syncytial virus. Science 342, 592–598 (2013).

66. J. P. M. Langedijk, et al., Universal paramyxovirus vaccine design by stabilizing regions involved in structural transformation of the fusion protein. Nat. Commun. 15, 4629 (2024).

67. B. Dadonaite, et al., Deep mutational scanning of H5 hemagglutinin to inform influenza virus surveillance. PLoS Biol. 22, e3002916 (2024).

68. T. J. C. Tan, et al., High-throughput identification of prefusion-stabilizing mutations in SARS-CoV-2 spike. Nat. Commun. 14, 2003 (2023).

69. P. Schommers, et al., Restriction of HIV-1 escape by a highly broad and potent neutralizing antibody. Cell 180, 471–489.e22 (2020).

70. C. E. Radford, J. D. Bloom, Comprehensive maps of escape mutations from antibodies 10-1074 and 3BNC117 for Envs from two divergent HIV strains. J. Virol. 99, e0019525 (2025).

71. M. B. Doud, J. M. Lee, J. D. Bloom, How single mutations affect viral escape from broad and narrow antibodies to H1 influenza hemagglutinin. Nat. Commun. 9, 1386 (2018).

72. S. Iketani, et al., Viral entry properties required for fitness in humans are lost through rapid genomic change during viral isolation. MBio 9 (2018).

73. B. Sawatsky, D. A. Bente, M. Czub, V. von Messling, Morbillivirus and henipavirus attachment protein cytoplasmic domains differently affect protein expression, fusion support and particle assembly. J. Gen. Virol. 97, 1066–1076 (2016).

74. K. Voigt, et al., Fusogenicity of the Ghana Virus (Henipavirus: Ghanaian bat henipavirus) Fusion Protein is Controlled by the Cytoplasmic Domain of the Attachment Glycoprotein. Viruses 11 (2019).

75. H. C. Aguilar, et al., Polybasic KKR motif in the cytoplasmic tail of Nipah virus fusion protein modulates membrane fusion by inside-out signaling. J. Virol. 81, 4520–4532 (2007).

76. M. Weis, A. Maisner, Nipah virus fusion protein: Importance of the cytoplasmic tail for endosomal trafficking and bioactivity. Eur. J. Cell Biol. 94, 316–322 (2015).

77. L. Witte, et al., Epistasis lowers the genetic barrier to SARS-CoV-2 neutralizing antibody escape. Nat. Commun. 14, 302 (2023).

78. A. Moulana, et al., Compensatory epistasis maintains ACE2 affinity in SARS-CoV-2 Omicron BA.1. Nat. Commun. 13, 7011 (2022).

79. L. E. MacConaill, et al., Unique, dual-indexed sequencing adapters with UMIs effectively eliminate index cross-talk and significantly improve sensitivity of massively parallel sequencing. BMC Genomics 19, 30 (2018).

80. K. H. D. Crawford, J. D. Bloom, alignparse: A Python package for parsing complex features from high- throughput long-read sequencing. J Open Source Softw 4 (2019).

81. H. K. Haddox, et al., Jointly modeling deep mutational scans identifies shifted mutational effects among SARS-CoV-2 spike homologs. bioRxivorg 2023.07.31.551037 (2023).

82. T. C. Yu, et al., A biophysical model of viral escape from polyclonal antibodies. Virus Evol 8, veac110 (2022).

83. K. Katoh, D. M. Standley, MAFFT multiple sequence alignment software version 7: improvements in performance and usability. Mol. Biol. Evol. 30, 772–780 (2013).

84. B. Q. Minh, et al., IQ-TREE 2: New Models and Efficient Methods for Phylogenetic Inference in the Genomic Era. Mol. Biol. Evol. 37, 1530–1534 (2020).

85. E. C. Meng, et al., UCSF ChimeraX: Tools for structure building and analysis. Protein Sci. 32, e4792 (2023).

86. J. Jumper, et al., Highly accurate protein structure prediction with AlphaFold. Nature 596, 583–589 (2021).

87. M. Mirdita, et al., ColabFold: making protein folding accessible to all. Nat. Methods 19, 679–682 (2022).

