## Supplemental Figures for "Functional and antigenic constraints on the Nipah virus fusion protein"

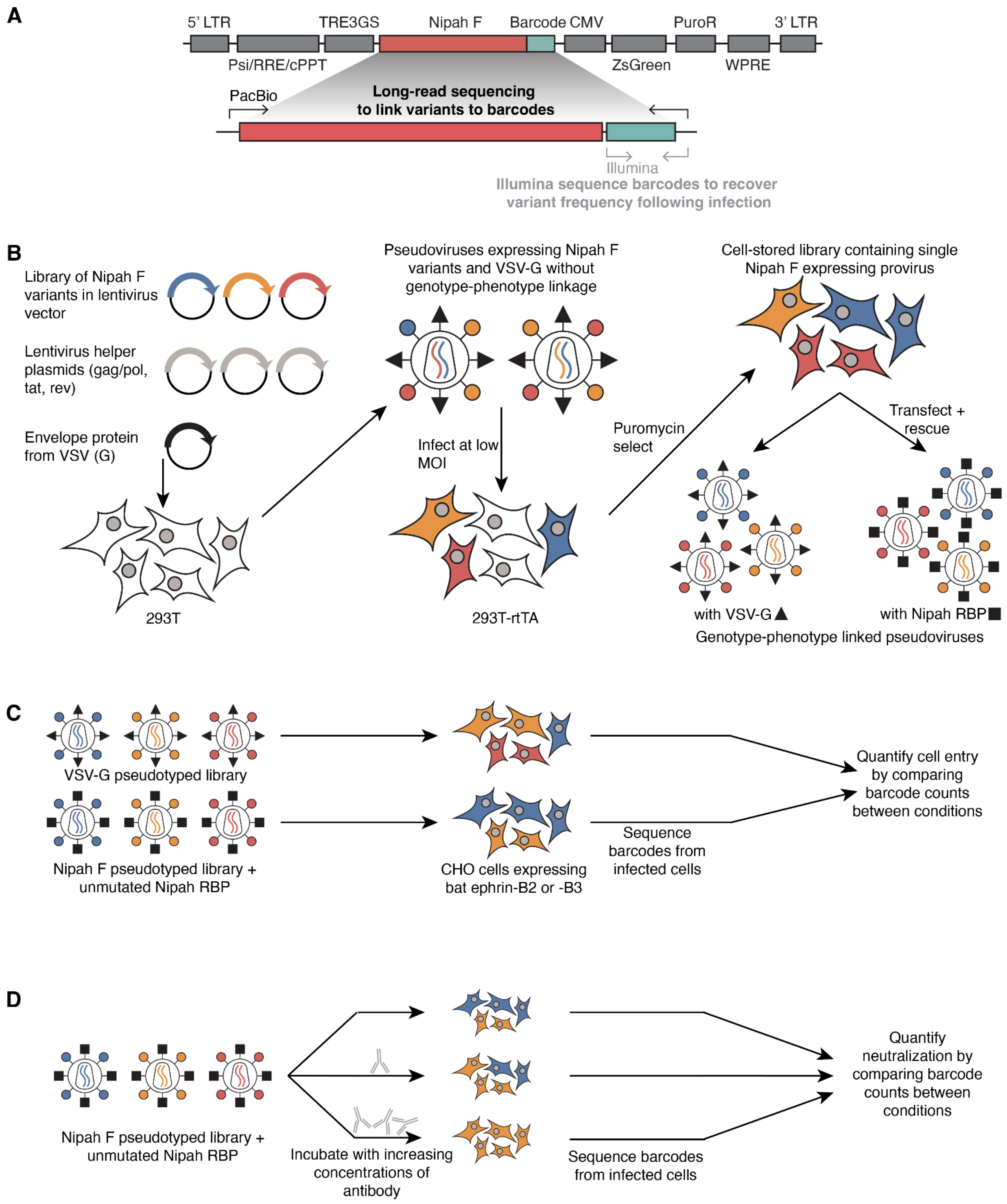


**Supplemental Figure 1. Graphical overview of the lentiviral backbone, production of lentivirus libraries, and selection strategies for generating deep mutational scanning data.**

**A)** Lentiviral backbone containing the Nipah F coding sequence (with a cytoplasmic tail truncation), essential lentiviral elements (long-terminal repeats, psi packaging element, rev response element, and central polypurine tract), and selection markers (ZsGreen and puromycin resistance). The F open reading frame is followed by a random unique 16-nucleotide barcode, which allows us to link specific F mutations with barcodes by long-read PacBio CCS sequencing. Downstream experiments rely on Illumina sequencing of the barcodes to obtain frequencies of the variants within each library. **B)** Creation of genotype-phenotype linked pseudovirus libraries. Lentivirus plasmid libraries containing all possible F amino acid mutations in the ectodomain (sites 29-481) are transfected into 293T cells with other lentiviral helper plasmids (gag/pol, rev, tat) and the envelope protein from the vesicular stomatitis virus (VSV-G), which has broad tropism. Pseudoviruses rescued from transfections cannot be used for deep mutational scanning due to mismatches between the genotype encoded in the pseudodiploid virion and the protein variants expressed on the surface. Instead, these pseudoviruses are used to infect 293T cells at a low multiplicity of infection (MOI < 0.01) to ensure a single integrated variant per cell, and bottlenecked to only get ~50,000-70,000 unique variants. Cells are passaged in the presence of puromycin, creating a cell-stored library of Nipah F variants contained within a lentiviral backbone. Pseudovirus libraries are rescued from these cells by re-transfecting essential lentiviral helper plasmids (gag/pol, tat, rev) and either VSV-G or the Nipah virus receptor binding protein. The final pseudovirus libraries express Nipah F variants with either VSV-G or the unmutated Nipah receptor binding protein. These libraries are used in the following steps to measure the effects of mutations. **C)** Overview of selection experiments used to obtain the effects of mutations on cell entry. Libraries generated in (B) that express Nipah F variants with either VSV-G or the unmutated Nipah virus receptor binding protein are used to infect stable CHO cells expressing either bat ephrin-B2 or -B3. Pseudoviruses with VSV-G on their surface will infect cells regardless of the F variant displayed and acts as a control for measuring the baseline composition of variants in the library. Pseudoviruses with the Nipah receptor binding protein and the F variants will only enter cells if they are functional. Twelve hours after infection, unintegrated lentiviral DNA templates are extracted from cells and undergo PCR and Illumina sequencing. Barcode frequencies are used to estimate the relative composition of the variant libraries in the different conditions and estimate the effects of specific mutations on cell entry. **D)** Overview of selection experiments used to obtain the effects of mutations on antibody neutralization. Pseudoviruses expressing the unmutated Nipah receptor binding protein and F variants are used to infect CHO cells expressing bat ephrin-B3. Pseudovirus libraries are either incubated with varying amounts of antibody or added directly to cells (as the control). Variant frequencies are obtained by extracting unintegrated lentiviral template DNA from the cells, amplifying the barcodes with PCR, and sequencing with Illumina. To obtain estimates of the amount of libraries that are neutralized, plasmid DNA containing known barcodes are spiked-in during DNA extraction.

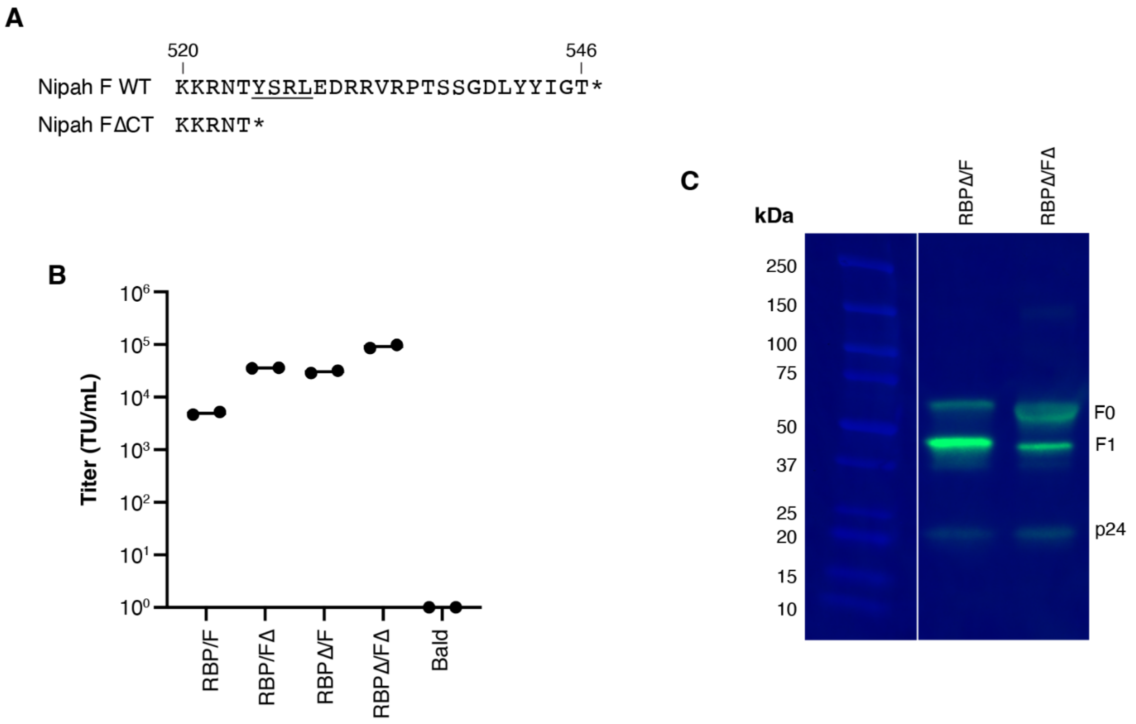


**Supplemental Figure 2. F cytoplasmic tail truncation affects pseudovirus titers and cleavage.**

**A)** Amino-acid alignment of wildtype (WT) and 22 amino-acid cytoplasmic tail deletion (ΔCT) at the C-terminus of Nipah virus F. Endocytosis motif 'YSRL' is underlined. **B)** Pseudovirus titers in CHO-bEFNB3cells of different combinations of RBP and F constructs with or without the cytoplasmic tail deletion. **C)** Reduced SDS-PAGE western blot on F and p24 in pseudoviruses (see Methods for details on staining). F0 and F1 are uncleaved and cleaved products, respectively. Gel is representative of two experiments.

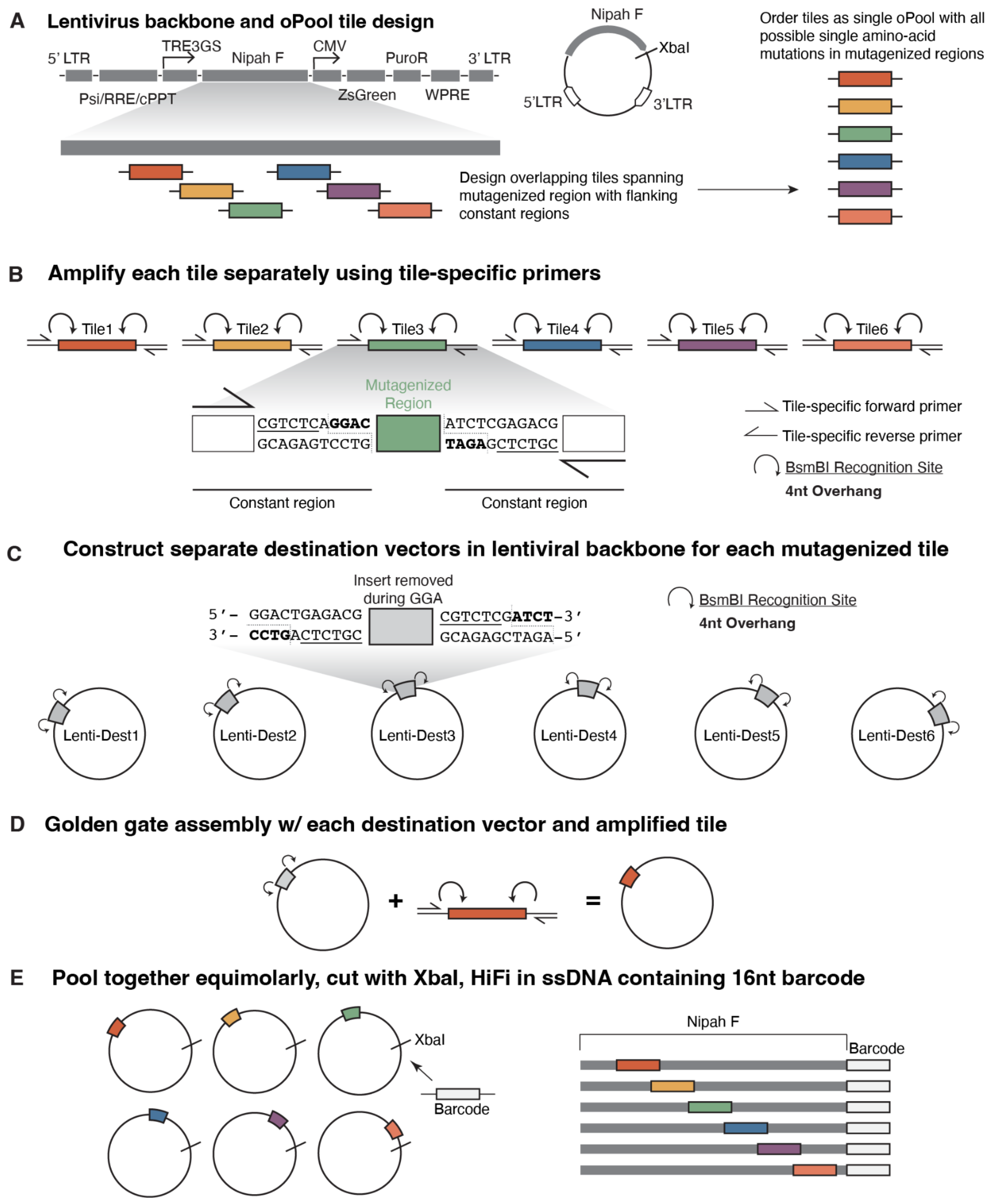


**Supplemental Figure 3. Schematic of single-mutation library construction using Twist oPools and Golden Gate Assembly.**

**A)** Plasmid map shows the different open reading frames of the parental F sequence contained downstream of a TRE3GS promoter within a lentiviral genome. The 3' LTR has been repaired so that lentiviral templates can be produced from provirus. Sites corresponding to amino acids 29-481 in Nipah virus F were divided into six overlapping tiles each spanning roughly ~220 nucleotides. Mutations to all possible amino-acid mutations at each position in the tile were constructed using the most frequent human codon. Finally, to facilitate downstream amplification of specific pools and cloning, constant regions were added to the 5' and 3' ends of each mutagenized window and contain BsmBI sites and unique forward and reverse primer sites. Each tile contains ~1500 mutations and was synthesized by Twist BioSciences. **B)** Each oPool tile was amplified with specific, unique primers that matched the unique priming site sequences included in the oPool design. **C)** We constructed six separate destination vectors based on the parental lentivirus genome in (A). We added inward-facing BsmBI sites at the flanks of each window using primer mutagenesis that result in 4 nt overhangs that match the overhangs in the tiles. **D)** Each amplified tile was mixed with the specific destination vector and assembled with Golden Gate Assembly. **E)** Once the tiles were cloned into each destination vector, we barcoded each plasmid. First, the six plasmids were mixed equimolarlly, followed by cutting with XbaI which is just downstream of the Nipah F ORF. HiFi reactions were done with the cut plasmids and a single stranded DNA oligo with flanks matching each side of the XbaI cut site and a 16nt random sequence in the middle.

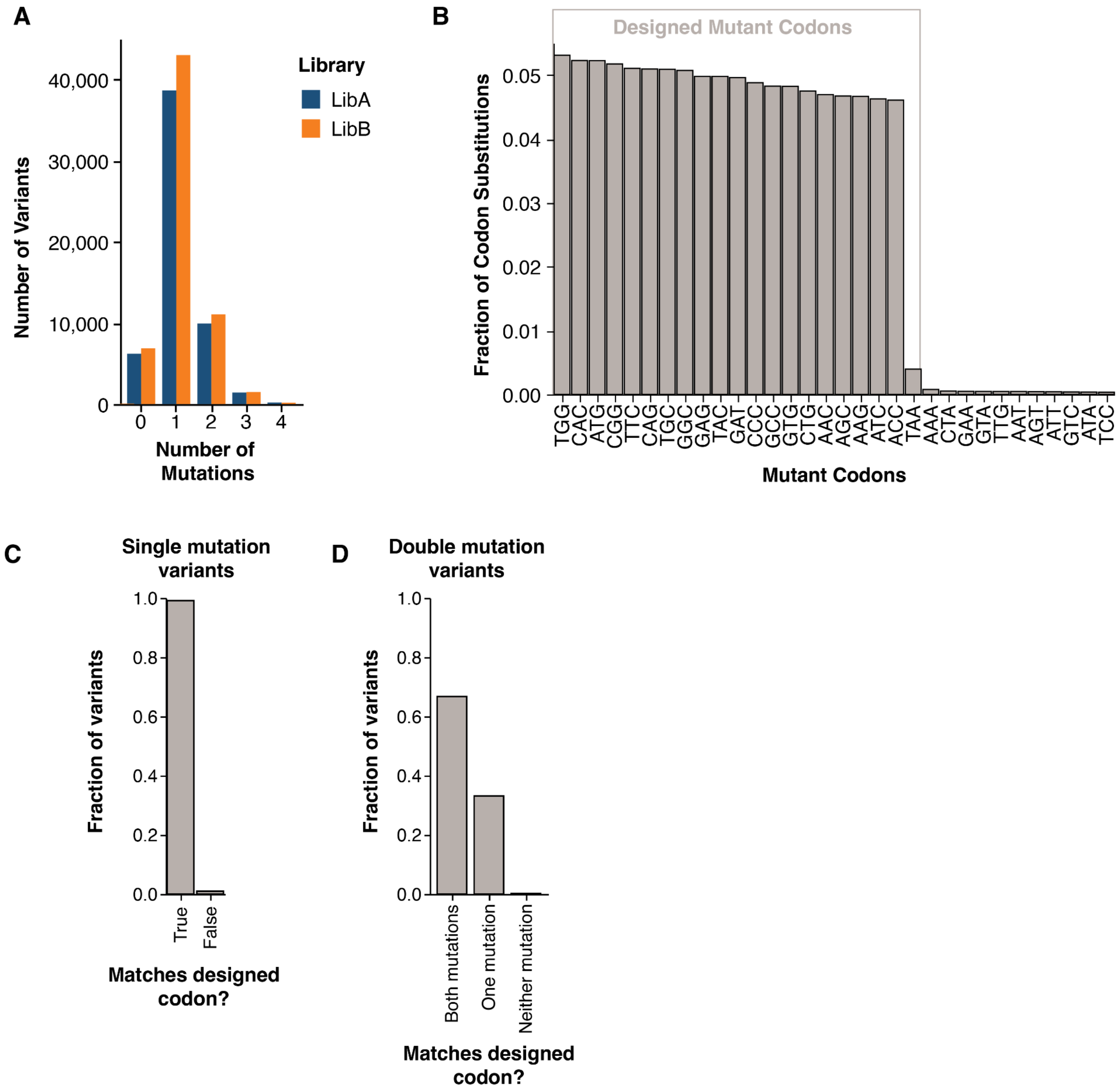


**Supplemental Figure 4. Deep mutational scanning library statistics.**
To link specific mutations with 16nt barcodes, we sequenced the full F open reading frame with PacBio along with the barcode downstream of the stop codon. Templates for sequencing were generated by rescuing pseudoviruses from cell-stored libraries expressing VSV-G and used to infect CHO cells. 12 hours later, unintegrated DNA viral templates were isolated and purified using a Qiagen Spin MiniPrep Kit, which were then used in PCR to generate amplicons for PacBio sequencing. **A)** Number of amino-acid mutations relative to the parental strain found in each variant. **B)** Frequency of mutated codons present in both libraries, with the specific codons used in oPool synthesis surrounded by a black box. There were 19 total missense codons at each site, and a subset of sites were mutated to a stop codon (TAA). **C)** Fraction of variants with one mutation relative to the parental strain that match the exact codon used for oPool design. **D)** Fraction of variants with two mutations that match the codons used for oPool

design. The majority of variants with two mutations match the codons used for oPool design at both sites, suggesting they are generated by lentiviral recombination rather than errors in oligo synthesis.

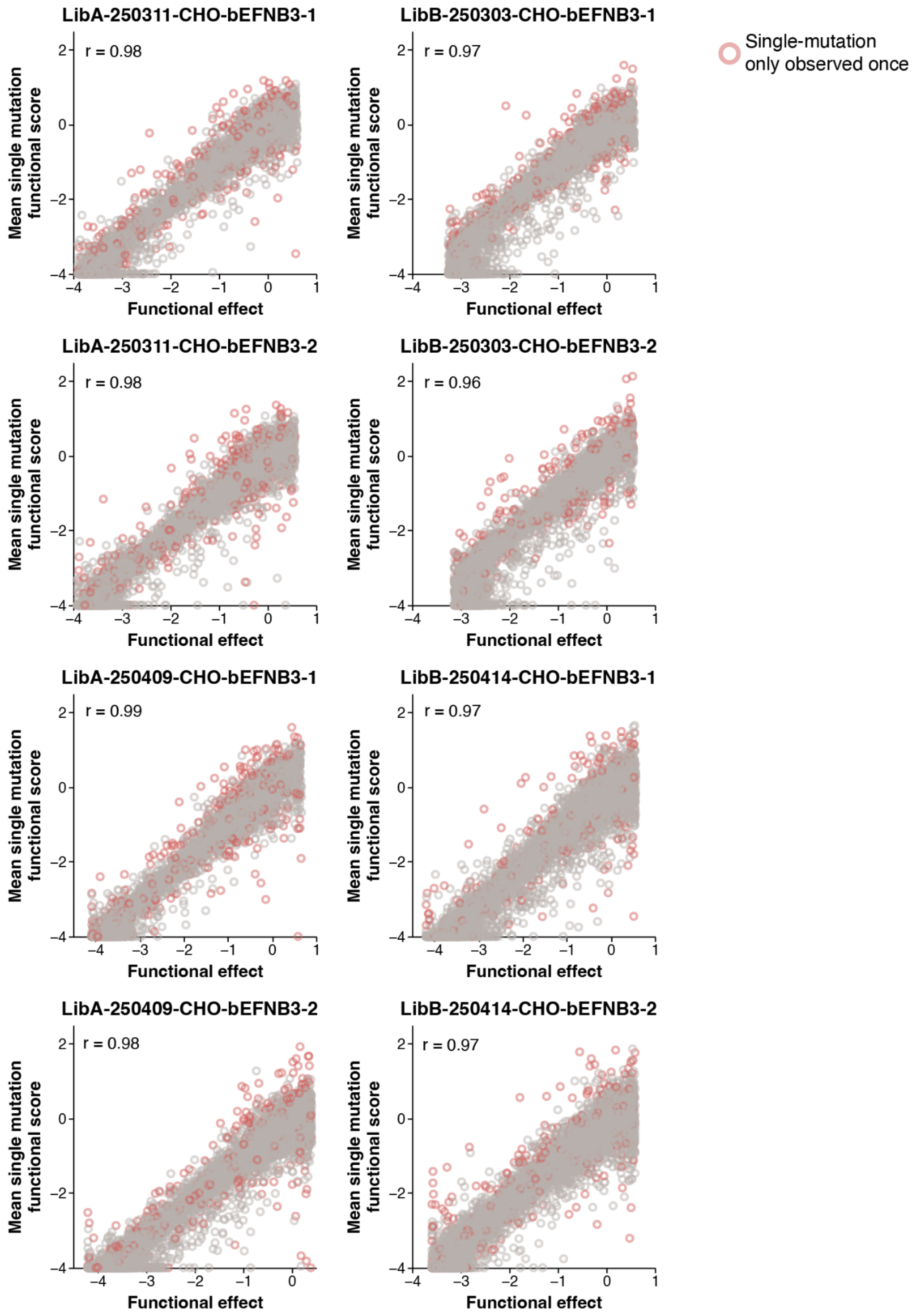


**Supplemental Figure 5. Correlation between functional scores from single-mutation variants and the decomposed functional effects following global epistasis fitting for four replicate selections each from LibA and LibB.**

Functional scores for each variant are calculated as the difference in frequency between the VSV-G and RBP/F pseudovirus infections. Functional effects are calculated by fitting a global epistasis model to decompose the effects of multiple mutations (49). Each plot shows an independent selection in CHO-bEFNB3 cells, and the r value is the Pearson correlation coefficient. Red points are mutations that are only linked to a single barcode and represent low-confidence measurements. Data were clipped at -4, corresponding to the lower range of detection for our assay.

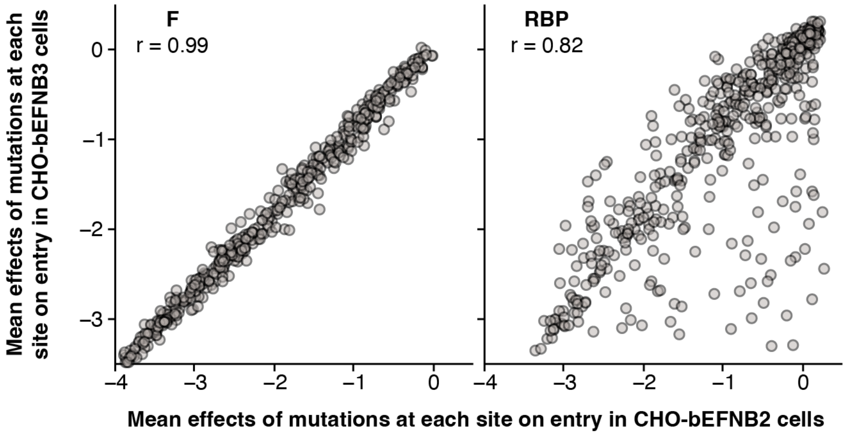


**Supplemental Figure 6. Mean effects of F and RBP mutations at each site on cell entry in CHO-bEFNB2 or -bEFNB3 cells.**

Correlation between the average effects of mutations on entry in CHO cells expressing either bEFNB2 or bEFNB3, with the measurements for F from the current study and RBP from Larsen et al. 2024 (41).

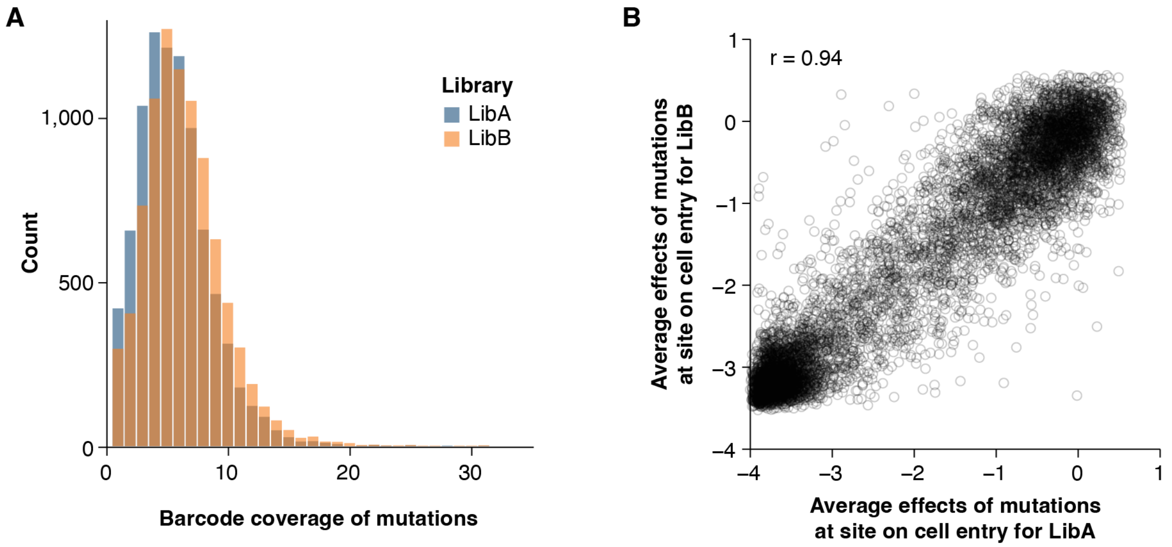


**Supplemental Figure 7. Barcode coverage of variants and correlation between independent replicate libraries.**

**A)** Number of unique barcodes covering each mutation following global epistasis fitting. Only mutations that were linked to at least two separate barcodes were included in the final analyses. **B)** Correlation in cell entry between replicate libraries. Average effects of mutations on cell entry were calculated separately for four independent selections with either LibA or LibB. These averaged effects were then used to estimate the correlation between each independent library.

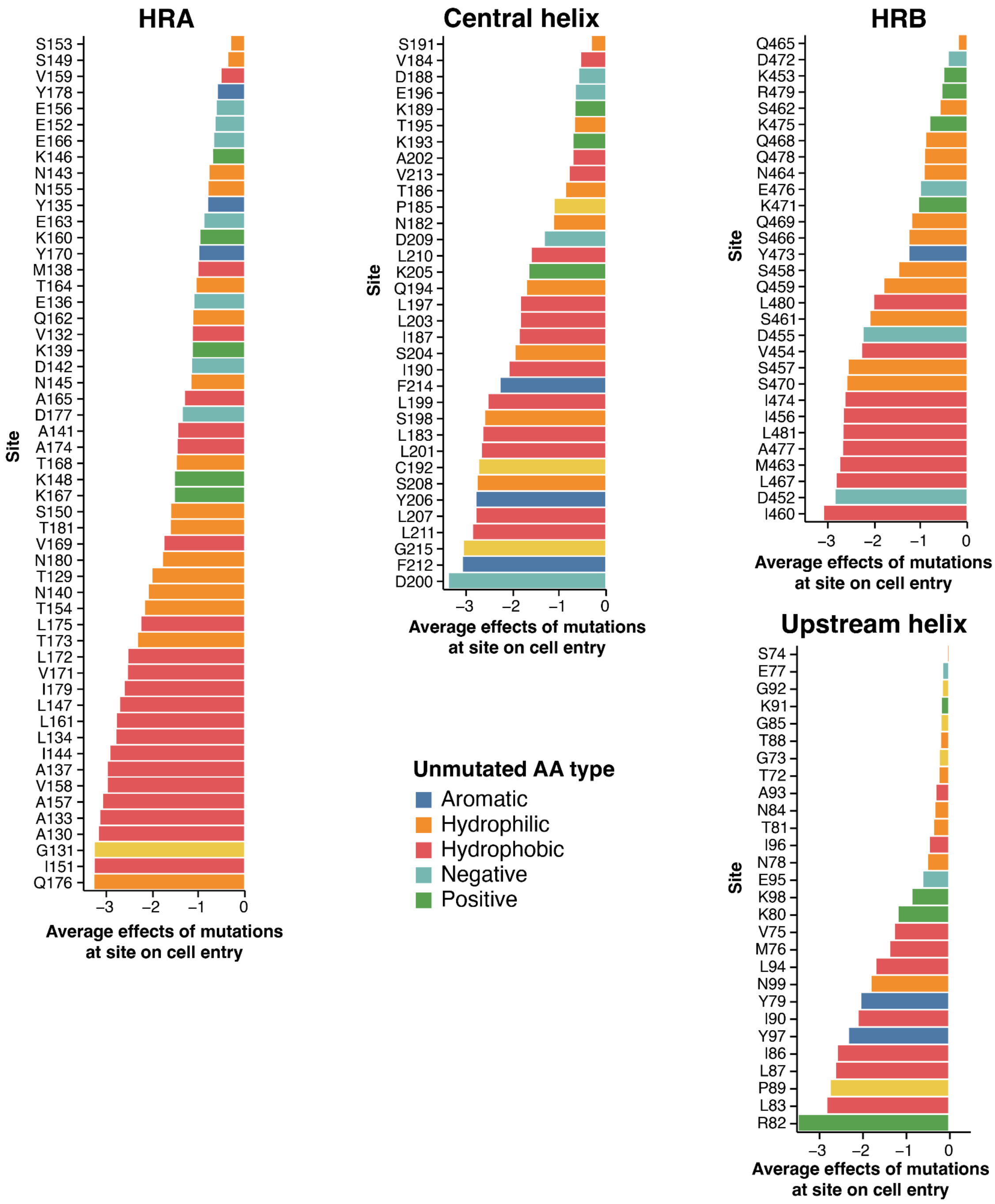


**Supplemental Figure 8. Average effects of mutations at each site on cell entry in the heptad repeat regions.**

Average effects of mutations at sites in the four main heptad repeat regions (Fig. 1). For each region, sites are ranked from least-constrained (top) to most-constrained (bottom) and are colored by the amino-acid property of the unmutated residue.

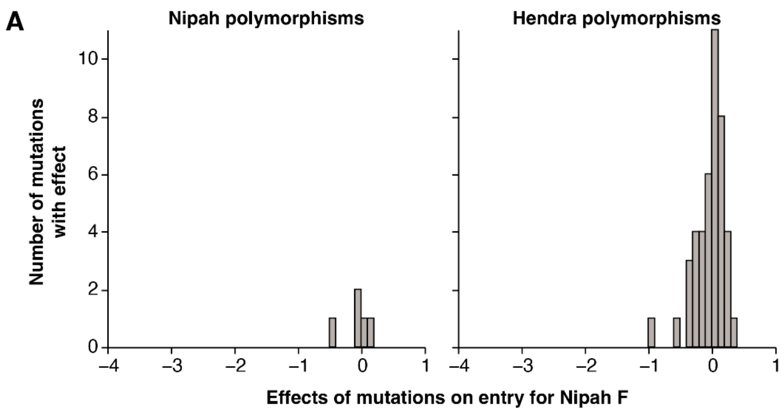


**B**

| Site | Parental Nipah AA | Mutation | # Observed | Cell Entry |
|---|---|---|---|---|
| 2 | V | A | 80 | |
| 6 | D | N | 80 | |
| 9 | C | Y | 80 | |
| 10 | Y | N | 2 | |
| 11 | C | S | 81 | |
| 13 | L | F | 2 | |
| 15 | I | L | 14 | |
| 19 | M | I | 21 | |
| 36 | I | V | 2 | -0.4 |
| 42 | V | I | 81 | 0.0 |
| 249 | A | T | 2 | 0.0 |
| 273 | S | G | 79 | 0.1 |
| 367 | L | M | 3 | -0.1 |
| 503 | A | V | 2 | |

**C**

| Site | Parental Nipah AA | Mutation | # Observed | Cell Entry |
|---|---|---|---|---|
| 42 | V | I | 16 | 0.0 |
| 68 | M | V | 20 | -0.1 |
| 70 | Q | K | 20 | 0.1 |
| 74 | S | T | 20 | -0.1 |
| 81 | T | S | 16 | 0.2 |
| 81 | T | N | 4 | 0.0 |
| 84 | N | T | 20 | -0.2 |
| 88 | T | S | 20 | 0.2 |
| 94 | L | I | 20 | 0.1 |
| 96 | I | L | 20 | -0.3 |
| 98 | K | N | 20 | -0.6 |
| 105 | V | I | 4 | -0.2 |
| 109 | R | K | 20 | 0.0 |
| 114 | I | V | 20 | 0.1 |
| 118 | V | I | 20 | 0.0 |
| 189 | K | Q | 20 | 0.2 |
| 198 | S | A | 19 | -0.1 |
| 267 | I | V | 20 | 0.2 |
| 293 | I | V | 20 | -0.3 |
| 316 | I | V | 20 | -0.3 |
| 318 | V | I | 20 | 0.2 |
| 328 | I | V | 20 | 0.4 |
| 329 | G | K | 20 | 0.0 |
| 330 | F | Y | 20 | -0.1 |
| 333 | I | T | 4 | 0.1 |
| 336 | R | K | 20 | 0.1 |
| 350 | N | A | 16 | 0.3 |
| 350 | N | S | 4 | 0.2 |
| 351 | N | S | 16 | 0.1 |
| 351 | N | T | 4 | -0.9 |
| 361 | E | D | 20 | -0.1 |
| 380 | N | G | 20 | 0.0 |
| 418 | P | T | 20 | 0.0 |
| 420 | A | V | 20 | 0.2 |
| 425 | V | I | 20 | 0.0 |
| 436 | V | I | 16 | -0.4 |
| 436 | V | T | 4 | -0.2 |
| 442 | G | S | 20 | 0.0 |
| 445 | I | V | 20 | 0.1 |
| 450 | F | Y | 20 | 0.0 |
| 453 | K | R | 4 | 0.2 |
| 479 | R | K | 16 | -0.1 |
| 480 | L | I | 20 | -0.3 |

**Supplemental Figure 9. Effects of mutations on cell entry in circulating Nipah virus and Hendra virus F sequences. A)** Distribution of measured effects of mutations in the F ectodomain on cell entry for all publicly available Nipah and Hendra F sequences. Each unique mutation relative to the parental Nipah F sequence was counted once. Only mutations found in at least two sequences are included. **B)** Information about each mutation found in circulating sequences relative to the parental Nipah F sequence. Although there were 14 mutations that occurred at least twice, most occurred in the signal peptide, which was not mutagenized for our experiments. **C)** Hendra F mutations in circulating sequences relative to the parental Nipah F sequence. Due to the large number of differences, only mutations that occurred in the ectodomain are shown.

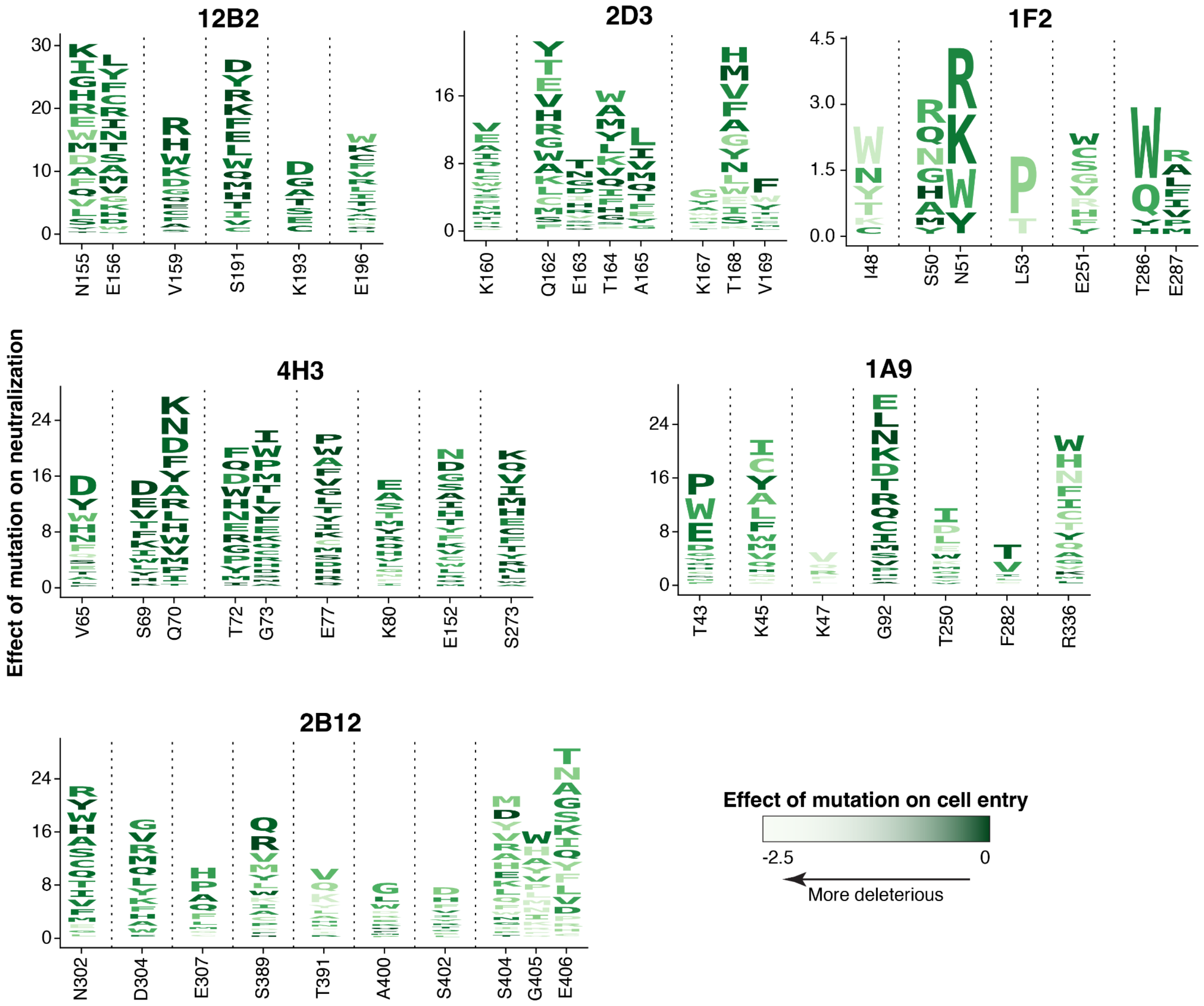


**Supplemental Figure 10. Effects of F mutations on antibody neutralization and cell entry.** Logo plots where the height of each letter is proportional to the amount a mutation decreases neutralization, and are colored by the effect that mutation has on cell entry, with darker colors indicating mutations that are neutral for cell entry, and lighter colors indicating mutations that are deleterious for cell entry. For each antibody, the logo plots show key sites where mutations have the greatest effect on neutralization.

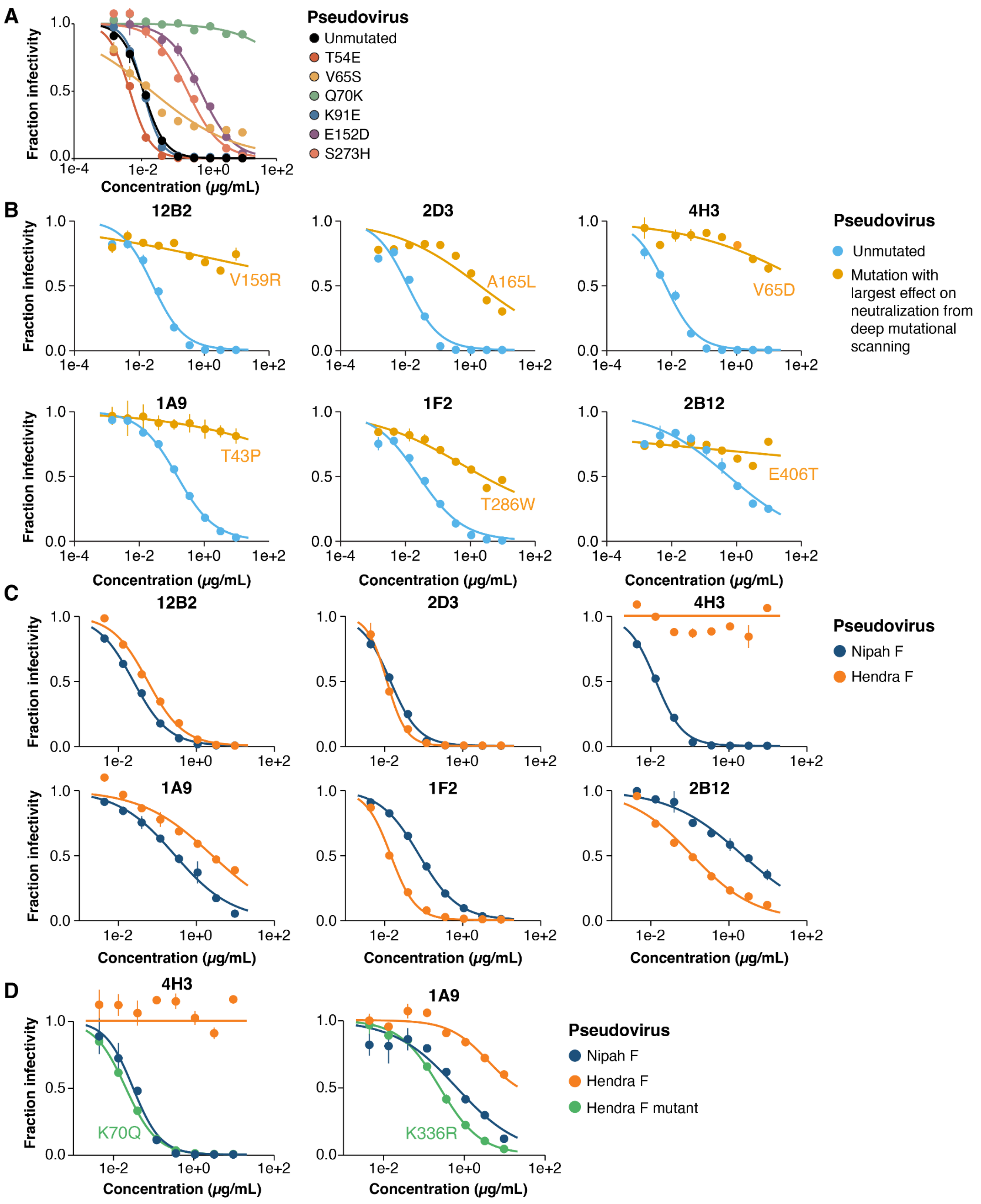


**Supplemental Figure 11. Neutralization curves by each antibody towards pseudoviruses expressing unmutated Nipah F, single-mutants of Nipah or Hendra F, or unmutated Hendra F.** Raw neutralization curves used for estimating $IC_{50}$ values for Figs. 6B, 6E, and 6G. Neutralization assays were performed using luciferase-based lentiviral particles pseudotyped with different Nipah F single mutations or Hendra F. Experiments were performed in duplicate. **A)** Neutralization curves of unmutated Nipah F and six different single mutations with the antibody 4H3. Individual mutations were selected to span a range of effects on neutralization. **B)** Neutralization curves of unmutated Nipah F and the mutation with the largest effect on neutralization for each antibody. **C)** Neutralization curves of unmutated Nipah and Hendra F. **D)** Neutralization curves of pseudoviruses expressing the unmutated Nipah receptor binding protein and either unmutated Hendra F, unmutated Nipah F, or Hendra F with specific mutations that revert the Hendra F to the amino-acid identity in the Nipah F at a site where this is expected to confer sensitivity to antibody neutralization.
